# Genotypic and phenotypic differences among phase-variable colony variants conserved across *Gardnerella* spp

**DOI:** 10.1101/2023.01.13.523976

**Authors:** Erin M. Garcia, Amy K. Klimowicz, Laahirie Edupuganti, Madeline A. Topf, Shraddha R. Bhide, Dawson J. Slusser, Samantha M. Leib, Gregory A. Buck, Kimberly K. Jefferson, Caitlin S. Pepperell, Joseph P. Dillard

## Abstract

The *Gardnerella* genus, now made up of more than 13 species, is associated with the polymicrobial disorder bacterial vaginosis (BV). However, the details of BV pathogenesis are poorly defined, and the contributions made by individual species are largely unknown. We report here that colony phenotypes characterized by size (large and small) and opacity (opaque and translucent) are phase variable and are conserved among all tested *Gardnerella* strains, representing at least ten different species. With the hypothesis that these different variants could be an important missing piece to the enigma of how BV develops in vivo, we characterized their differences. Beyond increased colony size, large colony variants (Lg) showed reduced vaginolysin secretion and faster growth rate relative to small colony variants (Sm). The ability to inhibit growth of *Neisseria gonorrhoeae* and commensal lactobacillus species varied by strain and in some instances differed between variants. Proteomics analyses indicate that 127-173 proteins are differentially expressed between variants. Further, whole genome sequencing analyses revealed an abundance of genes associated with variable poly-guanine tracts, implicating slipped strand mispairing in *Gardnerella* phase variation, and illuminating the potential for previously unrecognized variability within clonal populations. Collectively, these results suggest that colony variants may be primed to serve different roles in BV pathogenesis.

## Introduction

*Gardnerella* spp. are Gram-positive urogenital tract pathogens^1^. Numerous lines of evidence indicate that *Gardnerella* play a key role in bacterial vaginosis (BV), a disease with multiple adverse outcomes that ultimately reduces the barrier integrity of the vaginal epithelium and increases the susceptibility to other urogenital pathogens^2–6^. It is hypothesized that *Gardnerella* serves as a primary colonizer of the vaginal mucosa, acting to dampen the host immune response and establish the BV biofilm prerequisite to the arrival of secondary colonizers, such as *Fannyhessea vaginae* and *Sneathia* spp^7,8^. However, the mechanistic details of *Gardnerella* pathogenesis remain poorly characterized, partly due to the lack of methods for efficient directed mutagenesis in *Gardnerella* spp. and also due to the substantial genetic and phenotypic heterogeneity within the genus^9–12^.

*Gardnerella* taxonomy has varied since the discovery of the genus in the 1950s and continues to be a work in progress. Initially characterized as a singular species, later studies consistently supported the division of the *Gardnerella* genus into at least four distinct subgroups, distinguished by differences in the 16S rRNA V6 region or in the gene for the universal 60 kDa chaperonin protein (*cpn60*)^13,14^. Each subgroup exhibits unique genome sizes, GC content, and gene content in addition to notable phenotypic differences^10,15^. Most recently, a regrouping of the *Gardnerella* genus into multiple (13+) species was officially proposed following whole genome sequencing of 81 strains and MALDI-TOF MS analysis of 10 strains^16^. Subsequent reports indicate that representatives from multiple species are simultaneously present during BV and that the distribution of these representatives is heterogenous between individuals ^17–19^. While some connections have been made between the virulence potential of specific subgroups/species and their association with BV, the details of their unique contributions within the vaginal microbiome are currently unknown^18,20–22^. This lack of clarity in *Gardnerella* taxonomy has led to difficulties in interpreting the results of previous studies that did not differentiate the species well and further complicates the characterization of the intricacies of host pathogen interplay during BV.

While heterogeneity at the “community” level is now being explored in more detail, the potential for clonal heterogeneity in *Gardnerella* isolates has yet to be investigated. Population heterogeneity during infection is commonplace among diverse bacterial pathogens. Occurring either before or in response to host contact, this variation has important implications for infection outcomes as cell-to-cell differences can alter both how the immune system responds and whether or not the group as a whole persists. This diversity is advantageous, increasing the odds of survival within a given environment. Accordingly, there are multiple mechanisms to generate this population level of heterogeneity. Phase variation, wherein bacteria reversibly switch the production of a certain factor “ON” or “OFF,” is one strategy frequently employed by pathogenic organisms^23–28^. One of the most common mechanisms to achieve this variation involves slipped strand mispairing, resulting in the gain or loss of a short, repeated sequence during DNA replication. These repetitive sequences, either homo- or hetero-polymeric tracts of nucleotides, within or upstream of a gene, ultimately alter the reading frame or lead to altered transcription or translation of a gene or operon^29^. This ability to “flip” a molecular switch helps bacteria to both respond rapidly to their changing environment and to also escape recognition by the host immune response^28^. For example, in *Neisseria gonorrhoeae*, phase variation controls the production of Opa proteins. Gain or loss of a five base pair repeat in the coding sequence leads to translation of the full-length sequence proceeding or being terminated due to a frameshift^30^. The differential expression of these surface proteins, encoded by up to 11 different genes, results in different abilities of the bacteria to bind CEACAM proteins or heparan sulfate proteoglycan on host cells, and affects the ability to survive interactions with neutrophils ^31–33^. Differences in gross colony morphology (small vs large, rough vs smooth, opaque vs translucent) is often an indicator of a phase variable phenotype and has been used to distinguish factor-producing and factor-deficient variants in numerous bacterial species^34–36^. Variation of Opa protein expression in *N. gonorrhoeae* directs the assembly of bacterial cells into opaque (Opa+) or translucent (Opa-) colonies.

In this study we describe variations in colony morphology that are conserved among multiple *Gardnerella* species. Colony size and opacity are phase variable, which allowed for the ability to distinguish and purify colony variants for characterization. In vitro assays performed with variants demonstrate distinct phenotypes with respect to growth, virulence factor production, and interaction with other bacteria. Proteomics analysis identified differences in metabolic profiles, with 127-173 proteins differentially expressed between variants. Ultimately, the differences observed in variants suggest that each may be better suited to survive under different environmental conditions, and this may serve as a bet hedging strategy to promote population success in the host. We hypothesize that these different variants could be an important missing piece to the enigma of how BV develops in vivo. Further, whole genome sequencing analyses showed an abundance of genes associated with variable poly-guanine tracts, implicating slipped strand mispairing in *Gardnerella* phase variation, and illuminating the potential for previously unrecognized variability within clonal populations. Through continued characterization of phenotypic differences and the mechanisms that drive them, we will gain a better understanding of how vaginal dysbiosis emerges from a state of health and may identify novel targets to impede BV development. Addressing both clonal and community variability will be vital for rational design of improved BV therapeutics.

## Results

### *Gardnerella spp*. exhibit colony size variants that are phase variable

Three *Gardnerella* strains classified as distinct species by *cpn60* alignment were chosen for detailed characterization (**Fig. S1**). During routine passage of strains on BHIFF agar, we observed the presence of colony morphologies with distinct sizes. The first morphotype, “large colony variant” (Lg) consisted of round, raised, colonies that were ∼0.5-1.0 mm in diameter. The “small colony variant” (Sm) morphotype consisted of similarly shaped colonies that were significantly smaller in size (**Fig. 1, left**). On biofilm, sBHI, or HBT media, the difference in size between variants is not as notable. We also observed colonies with differences in opacity on sBHI medium (**Fig. 1, right**). Opaque (Op) colonies were white to off-white whereas translucent (Tr) colonies were gray to very nearly clear. While in our hands large colonies were predominantly also opaque and small colonies were predominantly also translucent, we have observed large, translucent and small, opaque colonies (data not shown). This observation suggests both phenotypes are independent of each other. We tested whether these observations were restricted to select strains or consistent across a variety of *Gardnerella* species and strains. In all fifteen strains tested, including representatives from ten different *Gardnerella* species, the presence of both colony morphologies was readily apparent (**Table S1**).

**Figure 1.**
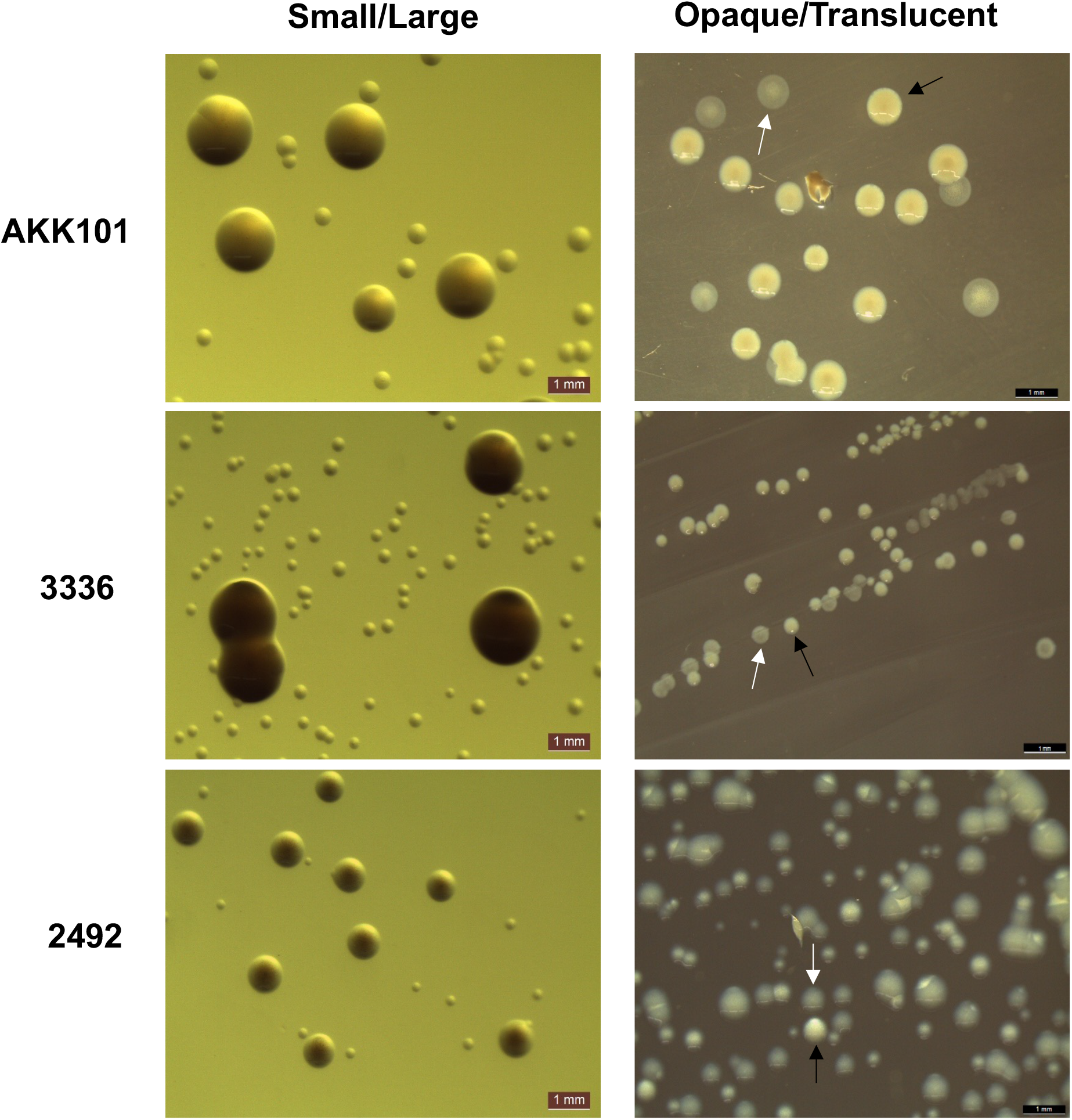
*Gardnerella spp*. exhibit variation in colony size and opacity. (Left) Strains from three different *Gardnerella* species assembled into large (Lg) and small (Sm) colonies on BHIFF agar medium. (Right) Opaque (Op) and translucent (Tr) colonies observed on sBHI plates. Black arrows indicate Op and white arrows indicate Tr. Images were taken with a Leica S8 APO stereo microscope.

For both size and opacity morphologies, we observed that the passage of single colonies consistently yielded a small fraction of the opposite variant (Lg gave rise to Sm, Tr gave rise to Op, etc.), indicative of phase variation. Quantification of switch frequencies for Lg and Sm phenotypes are presented in **Table 1**. These data suggest that colony size phenotypes are under control of high frequency phase variation. Switch dynamics were variable between strains, not only with respect to stability of a given phenotype (switch frequency), but also regarding the probability of switch in a given direction. For example, strain 2492 exhibited a lower switch frequency (∼10^−4^) and thus more stably maintained a given colony phenotype compared to strains AKK101 and 3336 with higher switch frequencies (∼10^−3^). While strains AKK101 and 2492 were more likely to switch from a Sm to a Lg than from a Lg to Sm, the opposite was true for strain 3336. From these observations we conclude that colony size in *Gardnerella* spp. is phase variable, which prompted us to test whether these variants were phenotypically distinct in other ways.

**Table 1.** Phenotypic switching of colony size occurs at appreciable frequencies in both directions. Data are presented as an average of three independent experiments. SD represents the standard deviation between independent experiments.

| Strain | Direction of switch | Avg. switch frequency | SD |
| --- | --- | --- | --- |
| AKK101 | Sm to Lg | 9.25E-03 | 1.29E-02 |
|  | Lg to Sm | 2.72E-03 | 3.95E-03 |
| 3336 | Sm to Lg | 1.16E-03 | 5.46E-04 |
|  | Lg to Sm | 2.26E-03 | 2.78E-03 |
| 2492 | Sm to Lg | 7.34E-04 | 1.15E-03 |
|  | Lg to Sm | 6.63E-04 | 8.01E-04 |

### Growth rate is faster for large colony variants than small colony variants

The appreciable difference in colony size between variants on certain media suggested that the Sm variant was growing more slowly than the Lg colony variant. To quantify the difference in growth rate, we performed growth curves in two liquid media: BHIFF and biofilm medium (**Fig. 2A** and **B**). Doubling time was then calculated from these curves (**Fig. 2C** and **Fig. S2**). In BHIFF medium, doubling times were ∼4-7X greater for Sm variants than for Lg variants in all tested strains. In biofilm medium, doubling time was not significantly different between Lg and Sm variants for any strain, consistent with what is observed during growth on biofilm agar. These data indicate that the differences in colony size for the variants reflect differences in growth of the bacteria and not just differences in bacterial interactions during growth on agar.

**Figure 2.**
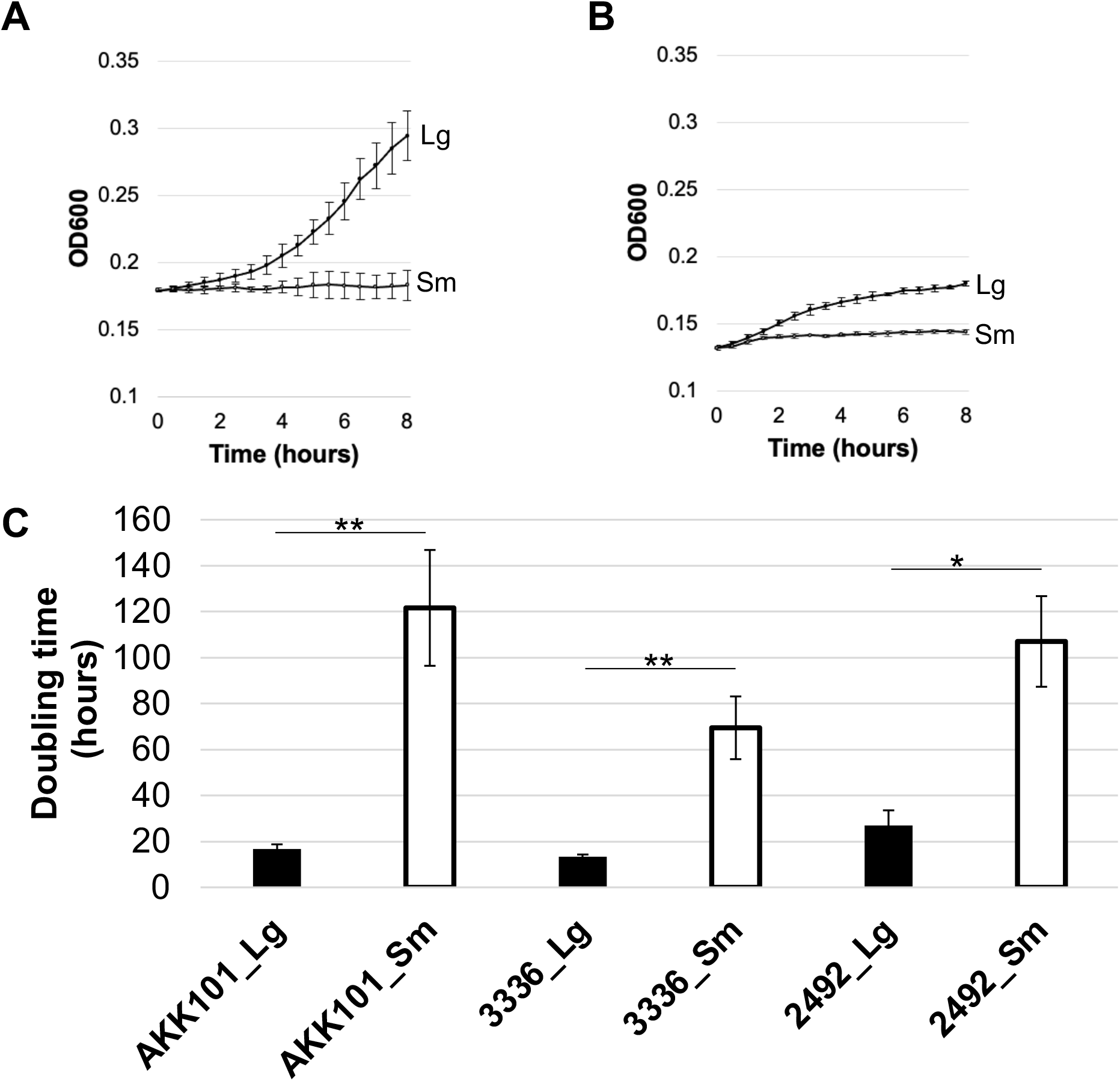
Large colony variants grow faster than small colony variants. Growth of each variant was quantified by measuring OD_600_ over a period of 8 hours. Curves are presented for *G. vaginalis* strain AKK101 growth in either BHIFF (A) or biofilm (B) medium. Data are presented as a representative experiment. Error bars represent the standard deviation among technical replicates. (C) Doubling time for each strain and variant in BHIFF medium. Error bars represent the standard error of the mean between at least three independent experiments. * p < 0.05, ** p < 0.01, *** P < 0.001.

### Small colony variants secrete greater amounts of vaginolysin than large variants

The small colony phenotype is associated with altered virulence factor production in some bacteria^37^. We sought to determine whether the secretion of vaginolysin (VLY), one of the most extensively studied virulence factors of *Gardnerella* spp., differed between variants. Strains were grown in biofilm medium, and the amount of VLY in culture supernatants at 24 hours post inoculation was assessed via hemolysis assays normalized to growth. The presence of VLY in culture supernatants was confirmed by Western blotting (**Fig. S3**). In all strains tested, hemolysis was significantly greater (5-19 fold) in small colony variants (**Fig. 4**). We also noted that hemolysis was substantially different between strains. These data indicate that the phase variants do not only differ in growth characteristics but also in virulence potential.

**Figure 4.**
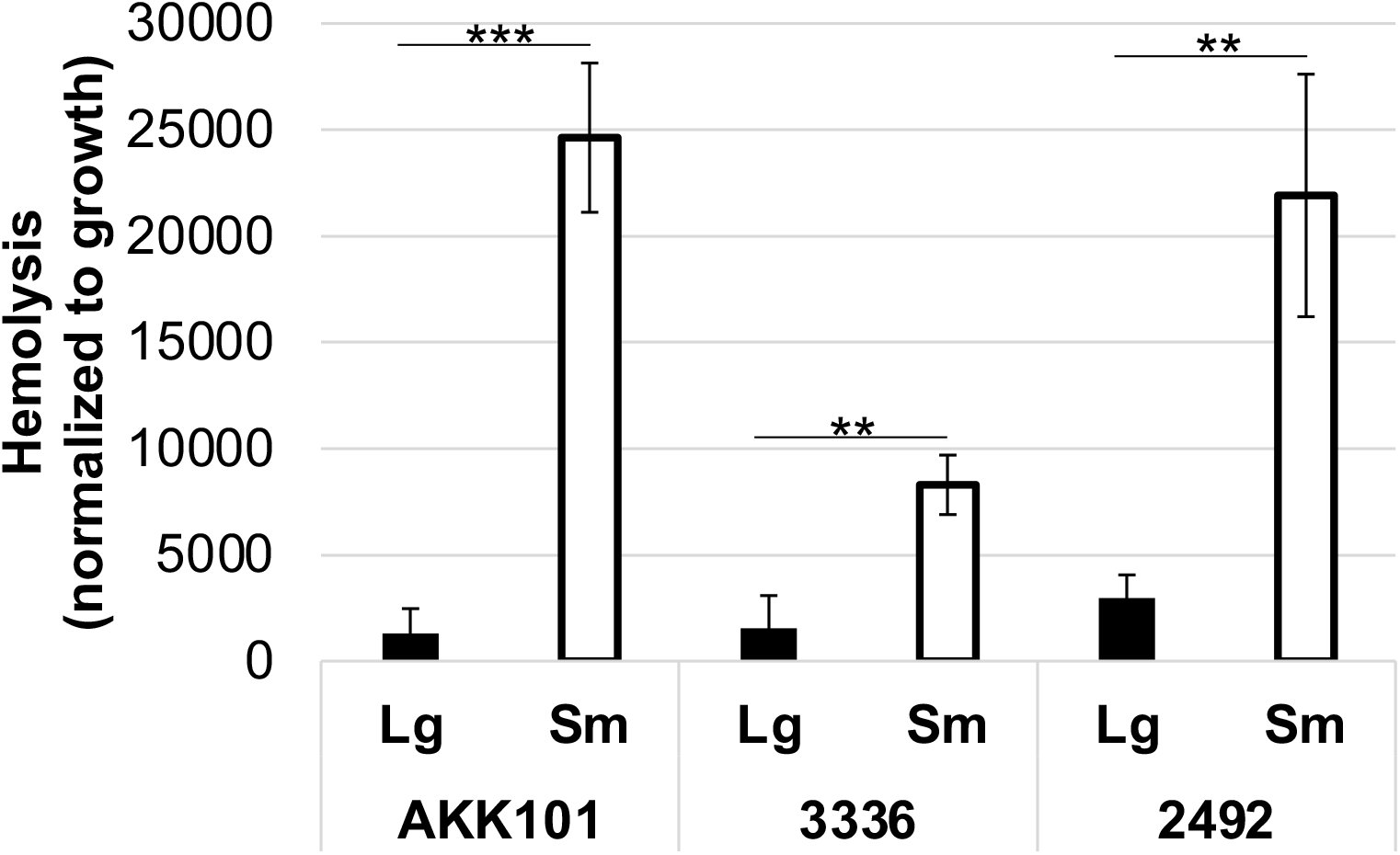
Small variant supernatants elicit greater hemolysis. Bacteria were grown in biofilm medium and the culture supernatant collected after 24 hours. Concentrated supernatant from triplicate wells was assayed for VLY production by percent hemolysis of human erythrocytes normalized to bacterial growth. Data are presented as an average of three independent experiments. Error bars represent standard deviation. ** p < 0.01, *** P < 0.001.

### Colony variants differ in their antagonism of vaginal commensals and pathogens

Multispecies interactions are important determinants of vaginal health and disease ^38,39^. We investigated the effects of *Gardnerella* variants from each species on the growth of representative commensal Lactobacillus species, *L. crispatus* and *L. gasseri*, and the urogenital pathogen, *N. gonorrhoeae*. These interactions are presented in **Fig. 5** and **Fig. S4**. The *Gardnerella* variants were streaked on a biofilm medium plate, and the following day a Lactobacillus or *N. gonorrhoeae* culture was spotted adjacent to each streak. None of the strains presented in this report affected growth of *L. gasseri*, determined by the absence of a zone of inhibition between the *Gardnerella* streak and the culture spot. However, it is worth noting that strains not characterized in this study did inhibit *L. gasseri* growth and inhibition differed between variants (data not shown).

**Figure 5.**
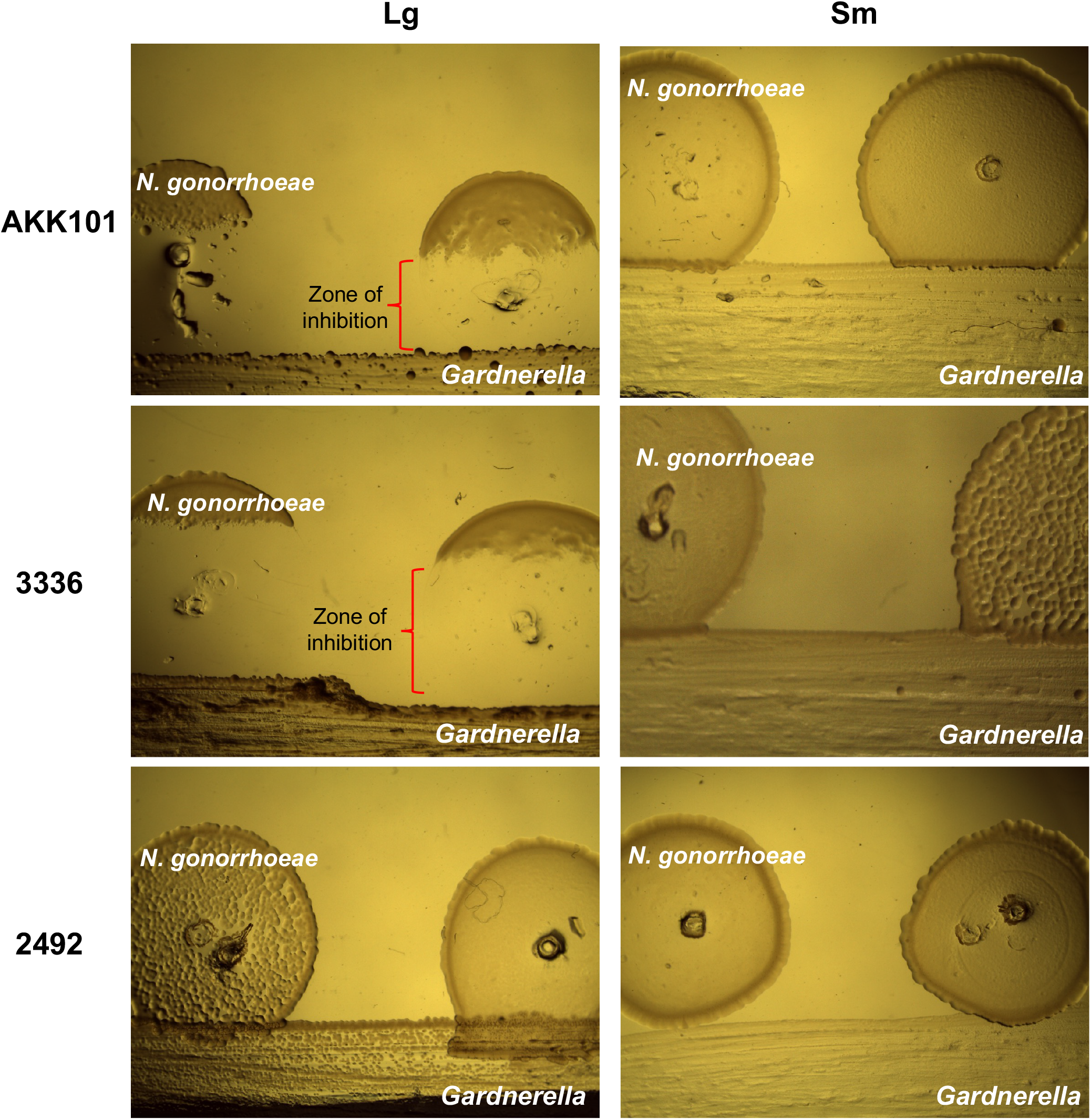
Colony variants differ in their antagonism of N. gonorrhoeae. *Gardnerella* colony variants were streaked onto biofilm medium + 10% FBS and grown for 24 hours before traversing the streak with 5µl of diluted *N. gonorrhoeae* cultures. Results from a representative experiment are shown but three independent experiments were performed. Images were taken on a Leica S8 APO stereo microscope.

In contrast to the results for *L. gasseri*, all of the strains characterized in this study exhibited some inhibition of *L. crispatus* growth, determined by the presence of a zone of inhibition with reduced *L. crispatus* density adjacent to the *Gardnerella* streak (**Fig. S4**). For strains AKK101 and 3336, this inhibition was observed for both Lg and Sm variants. Although, the zone of inhibition was significantly smaller for Sm variants. For strain 2492, inhibition was only observed for the Lg variant. The inhibition of *N. gonorrhoeae* by *Gardnerella* variants was substantially different from that of *L. crispatus*. The Lg variant of strains AKK101 and 3336 strongly inhibited *N. gonorrhoeae* growth, showing a large, complete (no single colonies) zone of inhibition. The Sm variant of these strains did not affect *N. gonorrhoeae* growth. Strain 2492 did not inhibit *N. gonorrhoeae*. These results demonstrate that *Gardnerella* spp. variants from some strains either have different responses to different bacteria or produce products that are inhibitory to select organisms.

### Proteomic differences between Lg and Sm variants

To begin to dissect the factors contributing to the many observed phenotypic differences in colony variants, we performed a proteomics analysis on each variant from strains of two different species. ATCC 14018 was chosen for inclusion, rather than the other strains presented in this study because, at the time its genome had already been sequenced. All data presented are of proteins detected as significantly different (p < 0.05) between variants in analyses of three independent experiments. The proteins with increased expression in Lg variants in both strains are presented in **Table 2**. These proteins are predominantly involved in amino acid and protein synthesis. Interestingly, the aminotransferase, class I/II protein that is upregulated in both Lg variants appears to be a transcriptional regulator of the MocR family. The proteins with increased expression in Sm variants in both strains are presented in **Table 3**. Consistent with our data from **Fig. 4**, VLY expression was significantly increased in Sm variants relative to Lg, and this difference was greater in strain 2492 (95-fold) than in strain ATCC 14018 (3.3-fold). We also observed increased expression of proteins involved in glycogen breakdown (MalQ), phosphate transport (PstS, PhoU) oxygen tolerance (NoxE), nucleotide synthesis (HMPREF0421_20874, PyrH, NrdD, Gnd, PflB), cell wall synthesis (MurA, HMPREF0421_20394), DNA recombination (RecA, FtsK, HMPREF0421_20072) and RNA posttranscriptional modification (HMPREF0421_20849). In addition to the similarly expressed proteins presented in **Tables 2** and **3**, each strain displayed unique differences in protein expression between Lg and Sm variants. The top 15 proteins with the highest Lg/Sm ratio are presented for strains 2492 and ATCC 14018 in **Tables S2** and **S3**, respectively and the top 15 proteins with the highest Sm/Lg ratio are presented in **Tables S4** and **S5**. Lg variants in strain 2492 exhibited increased expression of a calcium-translocating ATPase (18.5-fold), a DNA topoisomerase (15.5-fold), ribonuclease J (4.8-fold), a C1-like peptidase (3.4-fold), and RpoA (2.4-fold). In strain ATCC 14018, Lg variants showed increased production of an internalin B repeat protein (43-fold), an alpha-L fucosidase (9.8-fold), ATP synthase domains (9.2 and 8.4-fold), and beta-galactosidase (4.7-fold). Of interest in strain 2492 Sm variants was the increased production of a ribosome hibernation promoting factor (9.9-fold), a predicted GA module surface protein (7-fold), and a universal stress family protein (4.7-fold). In Sm variants of ATCC 14018, expression levels of a glycogenase (50-fold), a GA module-containing protein (24-fold), and an arylsulfatase (10-fold) were significantly increased.

**Table 2.**
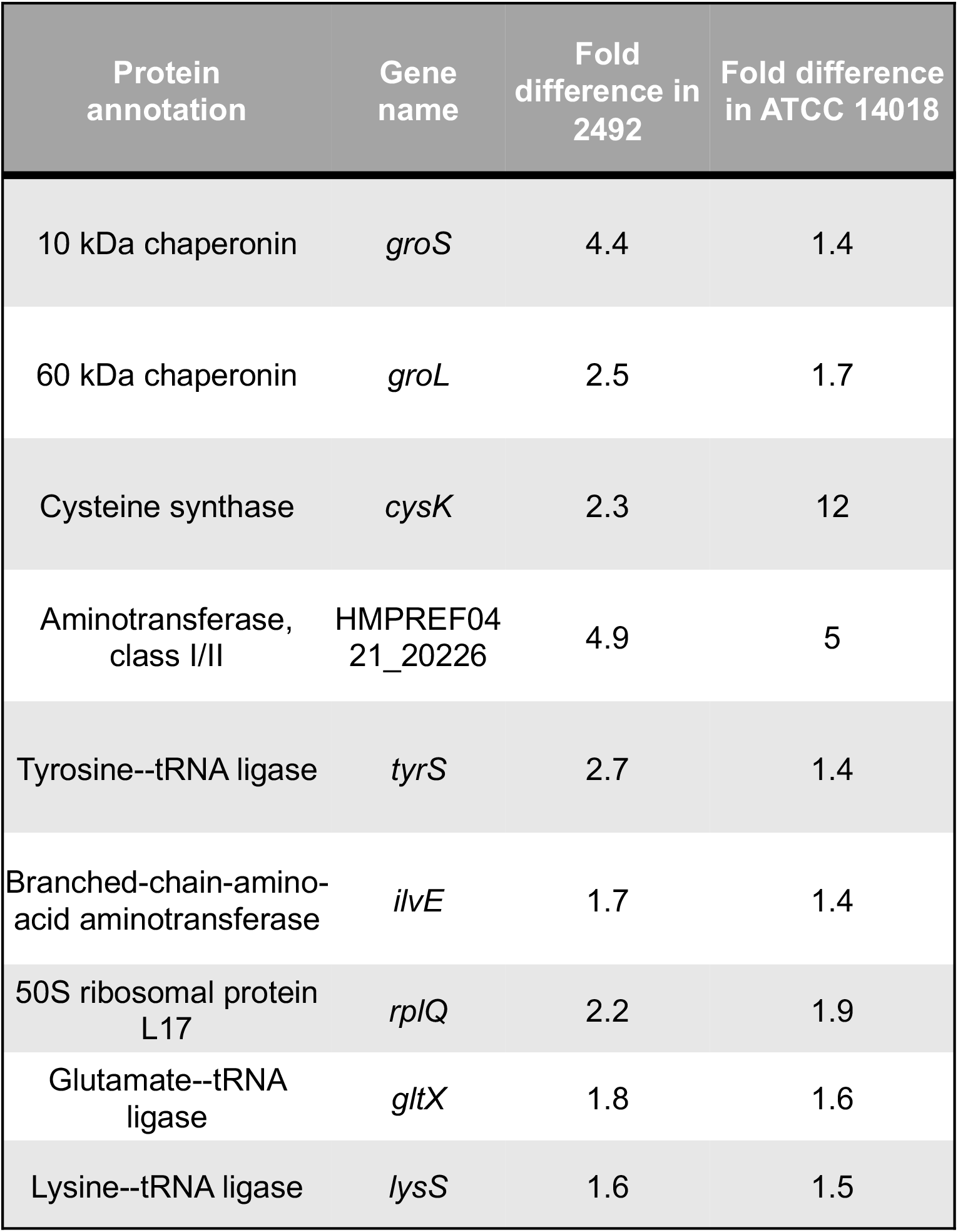
Proteins with greater expression in large variants. Proteins with similar expression pattern in both 2492 and ATCC 14018.

| Protein annotation | Gene name | Fold difference in 2492 | Fold difference in ATCC 14018 |
| --- | --- | --- | --- |
| 10 kDa chaperonin | *groS* | 4.4 | 1.4 |
| 60 kDa chaperonin | *groL* | 2.5 | 1.7 |
| Cysteine synthase | *cysK* | 2.3 | 12 |
| Aminotransferase, class I/II | HMPREF04<br>21_20226 | 4.9 | 5 |
| Tyrosine--tRNA ligase | *tyrS* | 2.7 | 1.4 |
| Branched-chain-amino-acid aminotransferase | *ilvE* | 1.7 | 1.4 |
| 50S ribosomal protein L17 | *rplQ* | 2.2 | 1.9 |
| Glutamate--tRNA ligase | *gltX* | 1.8 | 1.6 |
| Lysine--tRNA ligase | *lysS* | 1.6 | 1.5 |

**Table 3.**
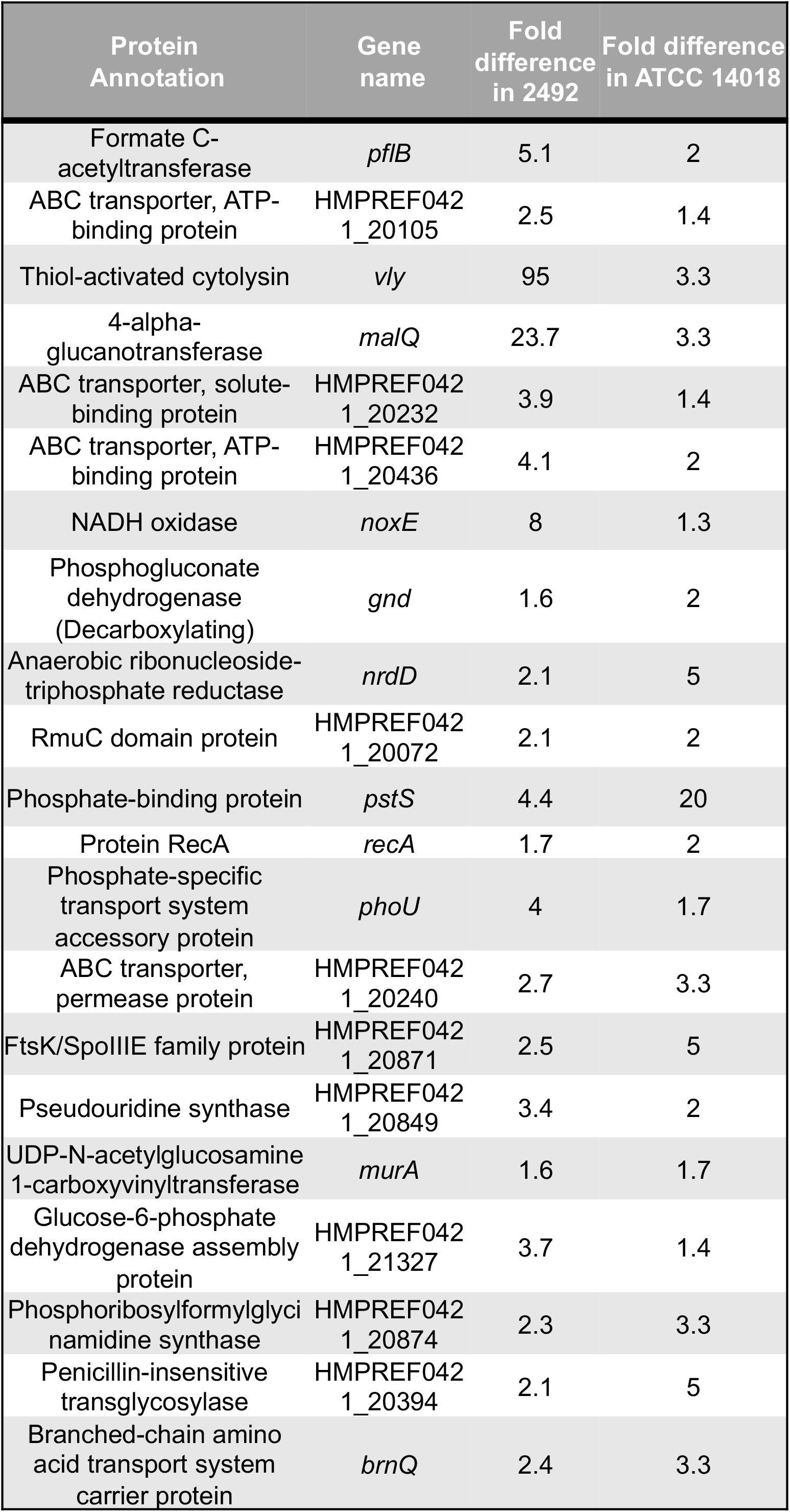
Proteins with greater expression in small variants. Proteins with similar expression pattern in both 2492 and ATCC 14018.

We next assigned all proteins with 1.5-fold or greater difference (p < 0.05) a KEGG protein class to determine if there were any functional differences between variants (**Fig. 6**). Consistent between both strains was the observation that a greater proportion of proteins were assigned to the “genetic information processing” class in Lg (inner circles) compared to Sm variants (outer circles). In Sm variants, the proportion of proteins left unclassified was greater than in Lg variants. Other functional differences observed were restricted to a single strain. For example, in ATCC 10418, the proportion of “energy metabolism” proteins was greater for the Lg variant while “transport” proteins were increased in the Sm variant. In 2492, “nucleotide metabolism” was increased in the Lg variant while “carbohydrate metabolism” was increased in the Sm variant. We also broke down the KEGG classes with the greatest number of protein assignments (transport, genetic information processing, amino acid metabolism, and carbohydrate metabolism) into subclasses to investigate whether there were any differences at this level (**Fig. S5**). For the “transport” class, peptide and nickel transport proteins were only observed in Lg variants. For “genetic information processing,” proteins classified into DNA replication, DNA repair, ribosome biogenesis, and chromosome and associated proteins subclasses were increased in Sm variants while proteins assigned to tRNA biogenesis, translation, and chaperones and folding catalysts subclasses were increased in Lg variants. The “amino acid metabolism” subclasses of glycine, serine, threonine metabolism and lysine biosynthesis were increased in Sm variants. Finally, the glycolysis/gluconeogenesis subclass of “carbohydrate metabolism” was increased in Sm variants.

**Figure 6.**
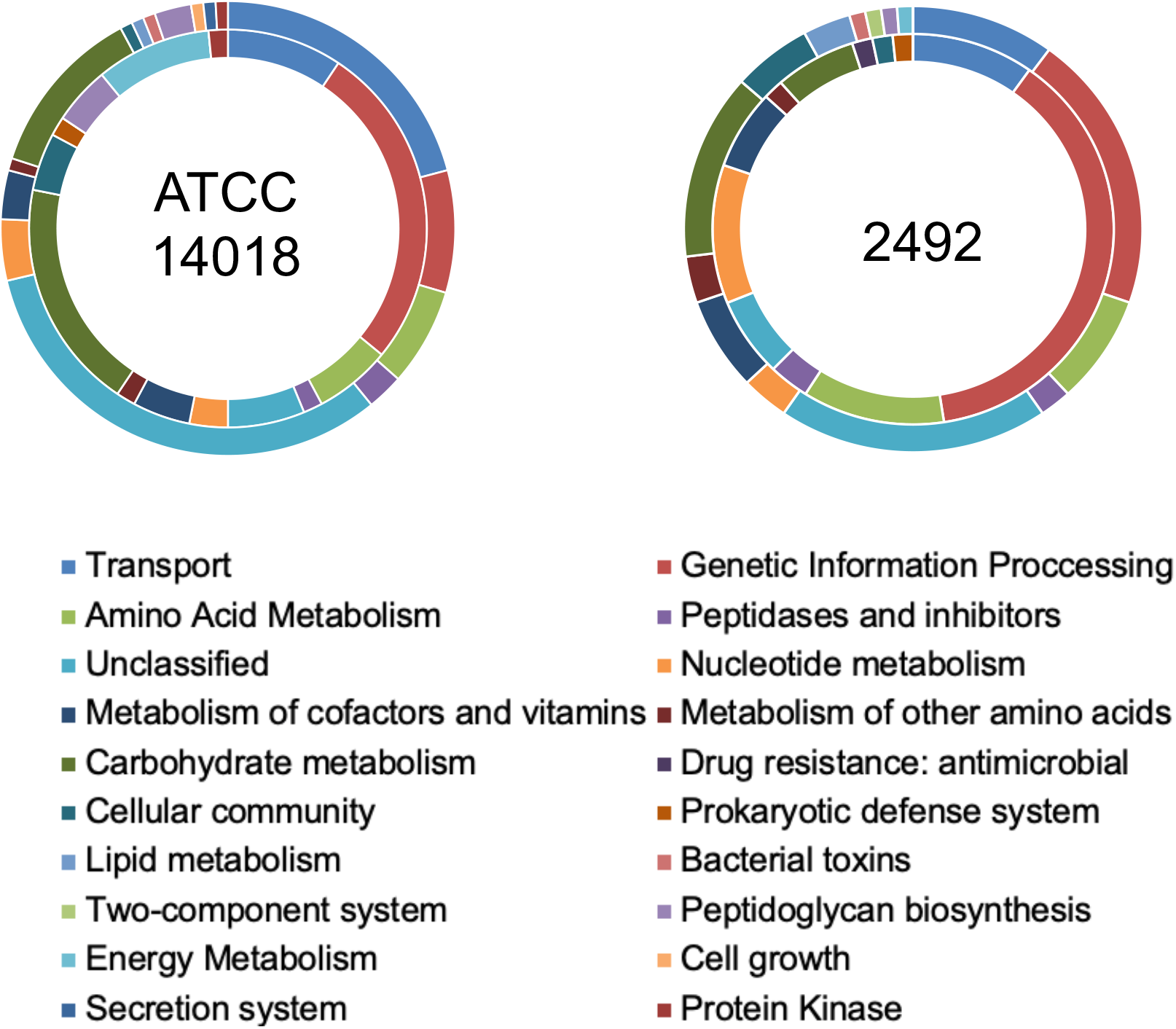
Protein classes differ by variant and strain. Proteomics data grouped by KEGG protein class. Inner ring is the Lg variant, outer ring is the Sm variant.

Collectively, these data demonstrate that Lg and Sm variants exhibit different metabolic profiles consistent with different growth characteristics and identify differentially expressed proteins likely to affect interactions with host cells. Furthermore, some differences between variants were found to be strain specific and may reflect the different roles of *Gardnerella* species in host colonization and disease.

### Poly-guanine tracts are abundant in *Gardnerella* spp. genomes

To define the genetic determinants of phase variation, we performed whole genome sequencing on variants from all strains of interest. Prior to genome comparisons between variants of the same strain for detection of SNPs and in/dels, we noted and characterized the ample distribution of homopolymeric guanine/cytosine tracts throughout all analyzed genomes (**Fig. 7A**). In each strain the average tract length was 10 base pairs. The majority of tracts (∼70%) were localized to putative promoter regions while the remainder, apart from one intergenic tract in strain 2492, were found within a predicted open reading frame. A majority were associated with ORFs annotated as hypothetical proteins, but these proteins commonly contained predicted Rib domains, GA modules, FIVAR domains, InlB B-repeat regions, and LPXTG motifs, suggesting localization to the cell surface. Further analysis of the number of homopolymeric guanine/cytosine tracts in publicly available *Gardnerella* genomes on NCBI (n=88) showed a median tract number of 20. This median differed when the strains were broken up by clade (**Fig. 7B**). Clade A, which includes *G. swidsinskii* and *G. leopoldii* species, had the lowest median at 18 tracts whereas clade C, which includes *G. vaginalis* and species group 2, had the highest median at 23 tracts. These results indicate that an abundance of genomic poly-guanine tracts is a conserved attribute of the *Gardnerella* genus. Due to the association of homopolymeric tracts with slipped strand mispairing and phase/antigenic variation, these results highlight a previously unrecognized potential for clonal heterogeneity among *Gardnerella* isolates.

**Figure 7.**
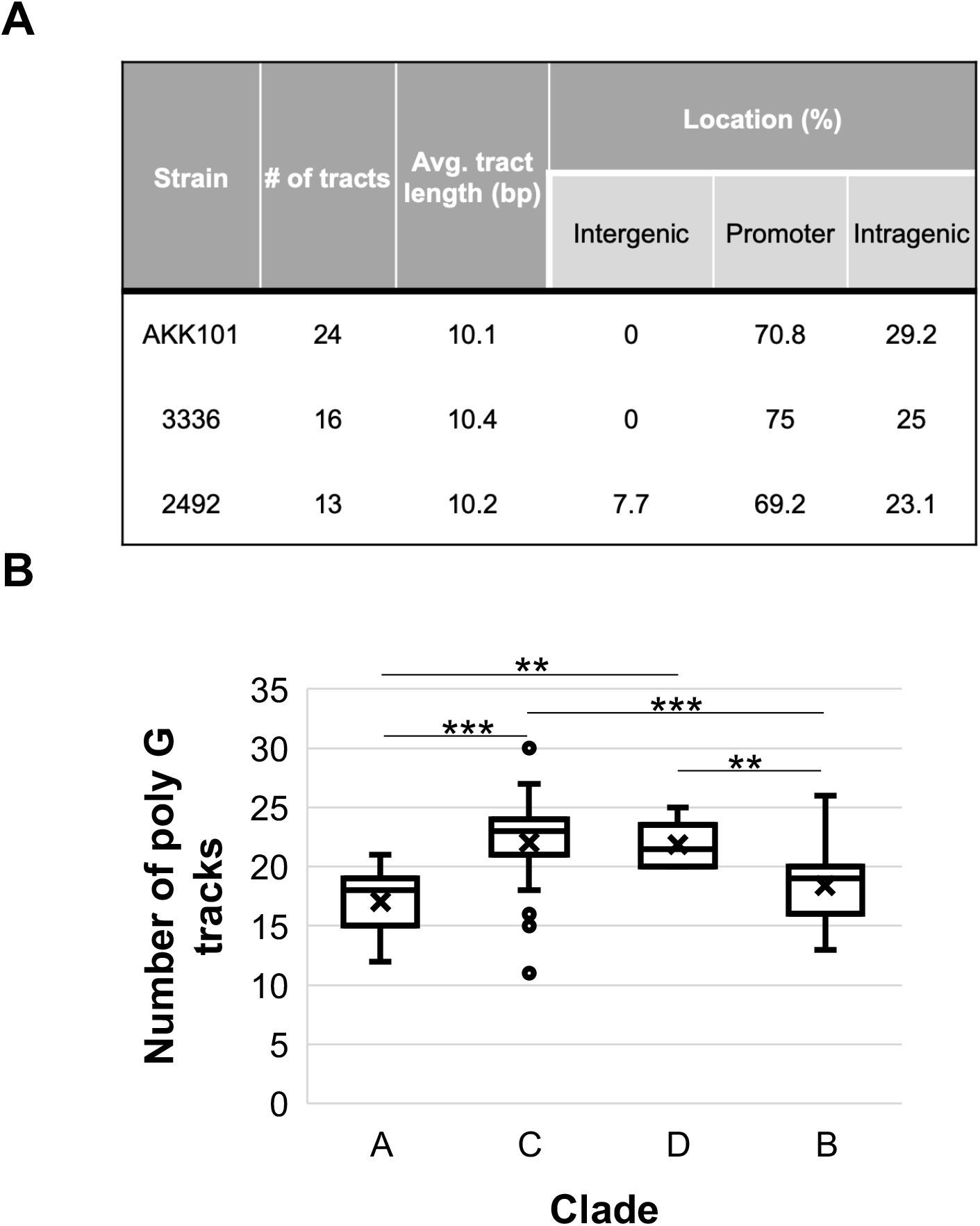
Poly guanine/cytosine tracts are abundant in *Gardnerella* genomes. (A) Characteristics of poly guanine/cytosine tracts in strains of interest. (B) Median tract number grouped by *Gardnerella cpn60* clade. ** p < 0.01, *** P < 0.001.

### Whole genome sequencing identifies multiple genotypic changes in *Gardnerella* colony variants

Mutation analysis called 45 single nucleotide polymorphisms (SNPs) and insertions/deletions (in/dels) for strain AKK101 and 35 for strain 3336. Between 26-38 % of variants called in each genome were within homopolymeric tracts. Missense (42-54%) and upstream variants (29%) were the most common mutations detected between colony variants, with frameshift mutations comprising a smaller but still appreciable portion (9-13%) (**Fig. 8**). Genes mutated in colony variants of both AKK101 and 3336 include *dps* (DNA protection from starvation), *ulaA* (ascorbate specific PTS transporter), and FIVAR and InlB B-repeat containing ORFs. Genes for ribosome proteins and tRNA ligases were also commonly mutated, but the specific gene varied by strain. These data support the idea that simple sequence repeats are a significant source of genotypic heterogeneity and identify candidate genes to be explored further for their role in colony phenotype.

**Figure 8.**
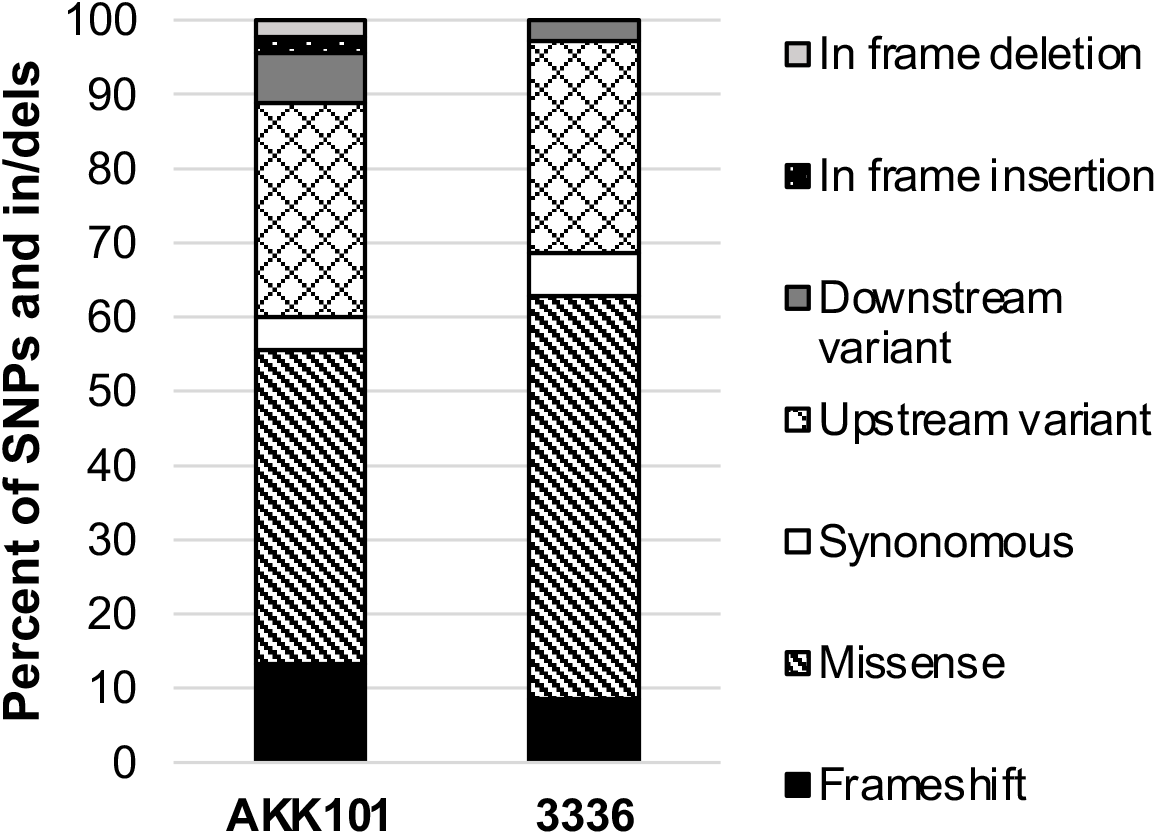
Missense mutations are most abundant between colony variants. Frequency of mutation classes called from comparisons of Lg and Sm variant genomes.

## Discussion

*Gardnerella* species are inhabitants of the lower human reproductive tract^40^. Though implicated in bacterial vaginosis, heterogeneity within the genus has hindered our understanding of the precise contribution(s) of each species. For many other pathogens, clonal heterogeneity imparted by phase and antigenic variation plays an important role in pathogenesis^41^. Yet, characterizations of these phenomena have been limited in *Gardnerella* spp^42^. In this study, we provide evidence for the existence of two phase variable colony morphotypes that were conserved across all tested *Gardnerella* isolates representing at least 10 different species. While colony differences among *Gardnerella* isolates were briefly noted in a report focusing on sialidase activity, the morphotypes were not characterized in detail^43^. We show here through in vitro assays and proteomics analysis that size variants are phenotypically distinct beyond their colony appearance, adding yet another layer of heterogeneity to *Gardnerella* spp. isolates. Further, whole genome comparisons simultaneously revealed an abundance of homopolymeric tracts within *Gardnerella* spp. genomes and provided evidence of their variation. Collectively, our study highlights the importance of colony phenotype and slipped strand mispairing to *Gardnerella* spp. pathogenesis.

In our characterization of colony size morphotypes, we observed that Lg colony variants displayed a faster growth rate and diminished cytotoxin production relative to Sm variants. The ability to inhibit growth of other bacterial species varied between morphotypes and strains. The proteomic profiles of variants were also distinct. While proteins involved in amino acid and protein synthesis were increased most (highest fold change) in Lg colonies, Sm colonies of both strains showed a more varied profile with increased representation of proteins related to glycogen breakdown, phosphate transport, oxygen tolerance, nucleotide synthesis, cell wall synthesis, DNA recombination, and RNA posttranscriptional modification. When all significantly increased proteins were grouped into functional categories, the proportion of proteins associated with genetic information processing was increased in Lg variants. Importantly, a majority of the proteomic differences between variants were strain and thus species specific. These data suggest two things. First, as has been observed by other groups, *Gardnerella* species are profoundly phenotypically distinct. Second, colony size variants exhibit distinct physiologies, suggesting each may be better suited to survive under different environmental conditions, and differentially produce factors that would likely impact their interactions within the host.

The distribution of *Gardnerella* spp colony phenotypes in vivo, and thus their role(s) in vaginal health and disease, is currently unknown. However, given the observed differences between Lg and Sm colony variants, we propose a few possibilities. First, it is possible that these different forms of the bacteria represent different stages in an infection cycle, similar to how *Chlamydia trachomatis* elementary bodies are the infectious but non-growing form of the bacteria^44^. In this scenario we hypothesize the Sm variant to be the initiator. Through slower growth kinetics and increased VLY production, it may be more resistant to the host innate immune response. Its enhanced production of glycogenases could promote the breakdown of glycogen into substrates able to be used by the Lg variant and potentially other BV-associated bacteria. In contrast, the faster growing Lg variant may be able to outcompete vaginal competitors for nutrient sources, form the biofilm mass that supports other BV-associated species, and support transmission via aggregates in vaginal secretions. This is supported by preliminary data showing increased biofilm formation in Lg variants (data not shown). Alternatively, it is possible that Sm variants represent a persistent form of the bacterium that functions to maintain infection following antibiotic or immune system challenges.

Through proteomic and genomic comparisons of multiple strains, we aimed to identify a causative factor in phase variable colony size. We reasoned that differences between variants detected in multiple strains would have an increased chance of contributing to phase variation, especially since the colony size phenotype was present in all tested isolates. We initially hypothesized a role for a single gene susceptible to slipped strand mispairing. However, the large number of changes in proteins accompanied by the ample distribution of variable homopolymeric tracts and strain to strain differences has complicated identification of such a factor. For strain 2492, the proteomics data provided some indication of mechanisms related to reduced growth/colony size. The Sm variant showed increased production of a ribosome hibernation factor and a universal stress protein, both associated with decreased metabolism^45,46^. However, these factors were not identified in the ATCC 14018 proteomic analysis, suggesting that different factors may be utilized in different species or that the differentially expressed proteins discussed above are not causative in determining growth characteristics. We are currently investigating the genomic hits shared between strains for their role in colony size and are analyzing strain 2492 for potential changes in DNA methylation.

Two ORFs associated with poly guanine tracts, putatively annotated as surface proteins, were called as variable between Lg and Sm variants in both strains. However, we believe it is more likely that they directly contribute to colony opacity, rather than size. That they were called in our analyses could simply be due to the fact that our stocks of Lg variants are predominantly Op and our Sm stocks are predominantly Tr. Nevertheless, we are currently pursing characterization of all homopolymeric tract-associated genes annotated as surface proteins for their association with colony opacity. We wonder if phase variation in *Gardnerella* spp. could be similar to that in *N. gonorrhoeae*, wherein 11 different genes encoding phase variable surface proteins at various loci throughout the genome can contribute in innumerable combinations to the colony phenotype^47^. This would be important to investigate further considering heterogeneity due to phase and antigenic variation has been a roadblock for gonococcal vaccine design for some time^48^. A comparable challenge may present itself should vaccine design for BV be pursued in the future.

In sum, the data in this manuscript present a novel account of *Gardnerella* heterogeneity. As suggested previously, Sm colonies may represent a developmental stage for the bacteria, facilitating particular steps in the infection process. If true, this would be a paradigm shift in our understanding of how *Gardnerella* contributes to the development of BV. The existence of a slower growing small colony variant may also explain the high rate of BV recurrence, as Sm colonies in other bacteria are associated with persistence and chronic disease states^49^. Quantifying the predominance of each variant *in vivo*, at various points during incident and recurrent BV, and testing whether certain environmental factors alter the predominance of each variant will be important next steps for hypothesis testing. Furthermore, phase variation of surface proteins in vivo could impede the immune response during BV, facilitating immune evasion to promote persistence and/or recurrence/re-infection. Identifying the repertoire of surface proteins susceptible to phase variation will be important for rational design of *Gardnerella*-targeting BV therapeutics.

## Methods

### Strains and culture conditions

*Gardnerella* spp. strains 3336 and 2492 were isolated from women attending the Seattle Sexually Transmitted Disease Clinic between 1978 and 1984. Strain AKK101 is a spontaneous erythromycin resistant derivative of strain Gv1, isolated from the University of Wisconsin-Madison in 2014. The sources of these strains and others referenced in the manuscript are outlined in the acknowledgements section. Various media were used for *Gardnerella* growth and variant distinction. These include HBT bilayer medium (Human Blood-Polysorbate 80, Thermo Scientific™), BHIFF (brain heart infusion (BHI) broth with 10% FBS and 10% Fildes Enrichment), sBHI (BHI with 1.0% [wt/vol] gelatin, 1.0% yeast extract, 0.1% starch, and 0.1% dextrose) broth or agar supplemented with 10% FBS, and biofilm (BHI with 0.5% yeast extract, 0.2% dextrose, 0.1% starch). The culture conditions used for a given experiment are detailed in each respective section below.

*Neisseria gonorrhoeae* strain MS11 was routinely cultured on GCB agar (Difco Laboratories) with Kellogg’s supplements I & II and incubated at 37°C with 5% CO_2_^50^ *Lactobacillus crispatus* 33820 (acquired from ATCC) and *Lactobacillus gasseri* were cultured on MRS agar at 37°C with 5% CO_2_.

### Clade and species determination

*Gardnerella* genomes (n = 91) were downloaded from NCBI and were annotated using Prokka^51^. Sequences from genomes that were annotated as “60 kDa chaperonin 1” by Prokka were used for constructing the phylogenetic tree. Phylogeny.fr was used to generate phylogenetic tree and ggtree was used to visualize and annotate the tree^52,53^. Clade and species designations were assigned manually based on the strategies used in Ahmed et al. 2012 (Clades 1-4), Jayaprakash et al. 2012 (Clades A-D), and Vaneechoutte et al. 2019 (13 species designations, 4 named)^54–56^.

### Growth rate determination

Colony variants were first inoculated into sBHI, grown for 24-28 hours anaerobically (5% CO_2_, 90% N_2_, 5% H_2_) and then then spun down and washed in PBS. Bacteria were then resuspended in growth media, adjusted to an initial OD_600_ of 0.1, and 100µl aliquots were added to a 96 well plate in triplicate. Bacteria were incubated at 37°C in a BioTek Synergy HT plate reader for 8 hours under aerobic conditions. Absorbance at OD_600_ was measured every 30 minutes. Growth curves were performed in two media types (BHIFF and biofilm) supplemented with 0.042% sodium bicarbonate. To determine the rate of growth, a linear segment of each growth curve was selected, and the equation of the fit line was used to calculate the time it would take for the OD_600_ to increase by two-fold.

### Switch frequency

Freezer stocks of colony variants were steaked onto HBT plates and incubated at 37°C with 5% CO_2_ for ∼24 hours. Growth was swabbed from plates into BHI broth, serially diluted, and plated onto BHIFF agar. Plates were incubated at 37°C with 5% CO_2_ for ∼48-72 hours. Colony phenotype was then assessed using a Leica S8 APO stereo microscope. Colonies (10-60) of each phenotype were collected on sterile filter paper and then added to BHI broth for serial dilution and plating. Plates were incubated at 37°C with 5% CO_2_ for ∼48-72 hours and the number of colonies of each phenotype were counted. Values for switch frequency represent the fraction of colonies that switched from the originally collected phenotype to the opposite phenotype.

### VLY hemolysis assay

*Gardnerella* strains were streaked onto HBT plates from freezer stocks and grown overnight at 37°C with 5% CO_2_. Growth was swabbed into biofilm medium and suspensions were adjusted to an OD_600_ of 0.1. Replicates of 200 µl of inoculum were placed in the wells of a 96-well tissue culture-treated plate and grown stationary at 37°C with 5% CO_2_ for 21-24 hr. The initial and final ODs were measured and culture supernatant from triplicate wells was combined. Cells were pelleted and 500 µl of supernatant was concentrated with a 30 kD cut-off column (Amicon Ultra 0.5 ml). Samples were stored at −80°C prior to analysis. Blood was obtained from healthy human donors according to a protocol approved by the University of Wisconsin-Madison Institutional Review Board. Erythrocytes were collected by centrifugation (500 x*g* for 5 min), plasma was removed, and the cells were washed three times with sterile PBS. A 1% solution of erythrocytes was prepared. Concentrated supernatant was serially diluted with PBS and 100 µl of each dilution was mixed with 100 µl of the 1% erythrocyte suspension in a V-bottomed 96-well plate. Positive and negative controls for hemolysis were a 0.1% solution of Triton X-100 (Tx) and PBS, respectively. The plate was incubated for 30 min at 37°C with 5% CO_2_ and then centrifuged for 10 min at 2000 rpm to pellet intact erythrocytes. Supernatant was transferred to a flat-bottomed 96-well plate and the absorbance (A) at OD_415_ was measured. Percent hemolysis was calculated as (A_sample_ – A_PBS_)/ (A_Tx_ – A_PBS_) X 100. These values were multiplied by the dilution factor and then normalized to the growth in biofilm media (difference between initial and final OD_600_).

### VLY Western blot

*Gardnerella* supernatants were prepared as outlined in the hemolysis section. Concentrated supernatants were mixed with SDS-loading dye and boiled for 5 min. Equivalent volumes of samples were run on 10% SDS-polyacrylamide gels, transferred to PVDF membranes (BioRad), and Western blots were performed essentially as described previously^57^. Primary VLY antibody 23A2 (Absolute Antibody) was used at a 1:10,000 dilution, and secondary goat anti-mouse HRP antibody (Santa Cruz Biotechnology) was used at a 1:20,000 dilution^58^. Millipore chemiluminescent HRP substrate was used per the manufacturer’s instructions. Membranes were imaged with the Licor Odyssey® Fc System. Stain UM035 was previously reported to lack the *vly* gene^59^.

### Bacterial antagonism

Freezer stocks of small and large colony variants were streaked onto HBT plates and incubated at 37°C with 5% CO_2_ for ∼24 hours. Growth was swabbed from plates into BHI broth and then 20ul was plated onto biofilm agar with 10% FBS via the drip plate method. Plates were incubated at 37°C with 5% CO_2_ for ∼24 hours. After 24 hours of growth, the *Gardnerella* streak was traversed with 5ul of serially diluted *N. gonorrhoeae, L. crispatus*, or *L. gasseri*. Dilutions of *N. gonorrhoeae and L. gasseri* were prepared from 24-hour growth on agar plates swabbed into BHI broth. *L. crispatus*, which did not consistently grow on MRS plates, was prepared from freezer stocks. Bacteria were co-incubated at 37°C with 5% CO_2_ for ∼24 hours before imaging on a Leica S8 APO stereo microscope.

### Preparation of whole cell lysates for proteomics analysis

Large and small colony variants of *Gardnerella* strains ATCC 14018 and 2492 were streaked on to HBT plates from freezer stocks and grown overnight at 37°C with 5% CO_2_. They were re-streaked onto biofilm plates containing 10% FBS and incubated for 1-2 days. The growth was swabbed into cold sterile PBS, and the cells pelleted at 4°C. Supernatant was removed and the cells were washed three times in cold PBS with final resuspension in 50mM sodium phosphate buffer (pH 7.5) with a 1X Roche protease inhibitor cocktail (2492) or a Pierce™ protease inhibitor tablet (ATCC 14018). The cells were lysed by sonication with a Branson digital sonicator (30 sec at 40% amplitude with a 1-sec on/off pulse), and insoluble material was removed by centrifugation (10 min at 13,000 rpm). Supernatant was transferred to a fresh tube and protein concentration was determined by Bradford assay. Samples were stored at −20°C prior to analysis. Three biological replicates were prepared for each strain and variant.

### Proteomics analysis

#### Enzymatic “in liquid” digestion

Protein aliquots (50µg) were incubated for 60 minutes on ice in 10% TCA and 33% (2492) or 50% (ATCC 14018) Acetone (final vol:vol) to precipitate proteins, spun for 10 minutes at room temperature with max speed (16,000xg), and then pellets were washed twice with cold acetone. Protein extracts were re-solubilized and denatured in 15 (2492) or 20 (ATCC 14018) μl of 8M urea in 50mM NH_4_HCO_3_ (pH 8.5) and then diluted to 60μl for reduction step. This dilution was in 2.5μl of 25mM DTT and 42.5μl of 25mM NH_4_HCO_3_, pH 8.5 for 2492 and 2.5μl of 25mM DTT, 5 μl of methanol and 32.5μl of 25mM NH_4_HCO_3_, pH 8.5, for ATCC 14018. Diluted extracts were incubated at 56°C for 15 minutes and cooled on ice to room temperature. Iodoacetaminde (3μl of 55mM) was added for alkylation and incubated in the dark at room temperature for 15 minutes. The reaction was quenched by adding 8μl of 25mM DTT. Finally, 8μl of Trypsin/LysC solution (100ng/μl 1:1 Trypsin (Promega) and LysC (FujiFilm) mixed in 25mM NH_4_HCO_3_) and 21μl of 25mM NH_4_HCO_3_ (pH 8.5) was added to 100µl final volume. Digestion proceeded for 2 hours at 42°C then an additional 4µl of trypsin/LysC mix was added and digestion continued overnight at 37°C. The reaction was terminated by acidification with 2.5% trifluoroacetic acid (TFA, 0.3% final concentration).

#### NanoLC-MS/MS

Digests were desalted using Agilent Bond Elute OMIX C18 SPE pipette tips per manufacturer protocol, eluted in 10µl of 60/40/0.1% ACN/H_2_O/TFA, and dried to completion in a speed-vac and finally. Samples were reconstituted in 40µl of 0.1% formic acid. Peptides were analyzed by nanoLC-MS/MS using the Agilent 1100 nanoflow system (Agilent) connected to a hybrid linear ion trap-orbitrap mass spectrometer (LTQ-Orbitrap Elite™, Thermo Fisher Scientific) equipped with an EASY-Spray™ electrospray source held at constant 35°C. Chromatography of peptides prior to mass spectral analysis was accomplished using a capillary emitter column (PepMap® C18, 3µM, 100Å, 150×0.075mm, Thermo Fisher Scientific) onto which 2µl of extracted peptides was automatically loaded. The NanoHPLC system delivered solvents A (0.1% (v/v) formic acid) and B (99.9% (v/v) acetonitrile, 0.1% (v/v) formic acid) at a rate of 0.50 µL/min to load the peptides over a 30 minute period and a rate of 0.3µl/min to elute peptides directly into the nano-electrospray. This was done with a gradual gradient from 0% (v/v) B to 30% (v/v) B over 150 minutes and concluded with a 10 minute fast gradient from 30% (v/v) B to 50% (v/v) B at which time a 7 minute flash-out from 50-95% (v/v) B took place. As peptides eluted from the HPLC-column/electrospray source, survey MS scans were acquired in the Orbitrap with a resolution of 120,000 followed by CID-type MS/MS fragmentation of 30 most intense peptides detected in the MS1 scan from 350 to 1800 m/z. Redundancy was limited by dynamic exclusion.

#### Data analysis

Elite acquired MS/MS data files were converted to mgf file format using MSConvert (ProteoWizard)^60^. For ATCC 14018, resulting mgf files were used to search against Uniprot *Gardnerella vaginalis* proteome databases (UP000001453, 10/2020 download, 1,365 total entries) along with a cRAP common lab contaminant database (116 total entries) while files for 2492 were used to search the *Gardnerella vaginalis* amino acid sequence database with a decoy reverse entries and a list of common contaminants (2,805 total entries). This was done using in-house *Mascot* search engine 2.7.0 (Matrix Science) with fixed cysteine carbamidomethylation and variable methionine oxidation plus asparagine or glutamine deamidation. Peptide mass tolerance was set at 15 ppm and fragment mass at 0.6 Da. Protein annotations, significance of identification and spectral based quantification was done with Scaffold software (version 4.11.0, Proteome Software Inc., Portland, OR). For ATCC 14018, peptide identifications were accepted if they could be established at greater than 94.0% probability to achieve an FDR less than 1.0% by the Scaffold Local FDR algorithm. Protein identifications were accepted if they could be established at greater than 98.0% probability to achieve an FDR less than 1.0% and contained at least 2 identified peptides. For 2492, peptide identifications were accepted if they could be established at greater than 97.0% probability to achieve an FDR less than 1.0% by the Scaffold Local FDR algorithm. Protein identifications were accepted if they could be established at greater than 99.0% probability to achieve an FDR less than 1.0% and contained at least 2 identified peptides. Protein probabilities were assigned by the Protein Prophet algorithm^61^. Proteins that contained similar peptides and could not be differentiated based on MS/MS analysis alone were grouped to satisfy the principles of parsimony. For ATCC 14018, proteins sharing significant peptide evidence were grouped into clusters.

KEGG protein category assignments were made manually by searching for gene names in the *Gardnerella vaginalis* ATCC 14019 genome (gvg) within the KEGG GENES database^62^. Proportions do not total to 1.0 as proteins could be classified into multiple categories.

### Genomic DNA extraction for sequencing

Strains AKK101 and 3336 were grown at 37°C anaerobically (5% CO_2_, 90% N_2_, 5% H_2_) on sBHI plates supplemented with 10% FBS. Bacterial pellets were lysed in colony lysis solution (1% Triton X-100, 2mM EDTA, 20mM Tris-HCL) with a “pinch” of lysozyme and the DNA was extracted using a DNeasy Blood & Tissue Kit (Qiagen). DNA quantity and quality were assessed via FEMTO Pulse at VCU Genomics Core.

### Whole genome sequencing

Whole-genome sequencing for strains AKK101 and 3336 was performed by the Virginia Commonwealth University Genomics Core using the MiSeq platform (Illumina, San Diego, CA, USA) after the construction of a paired-end sequencing library using the Illumina Nextera XT DNA sample preparation kit (Illumina, San Diego, CA, USA). Sequencing depth was 700X for AKK101 Lg, 600X for AKK101 Sm, 400X for 3336 Lg, and 200X for 3336 Sm.

### Sequencing analysis

Reads were assembled de novo using Shovill (https://github.com/tseemann/shovill) for AKK101 and 3336 and Flye for 2492^63^. Assemblies were annotated using Prokka and then large genomes were aligned against their respective small genome using BWA (https://github.com/lh3/bwa). SNPs were called using freebayse and SnpEff was used for annotating the mutations^64,65^. Poly guanine/cytosine tracts (8 or more consecutive nucleotides) were searched, counted, and characterized manually.

### Statistics

All data were assessed for normality using the Shapiro-Wilk test. Equality of variance was assessed using the Brown-Forsythe test. The means of normally distributed data with equal variance were compared by using a Student’s *t* test. The means of normally distributed data with unequal variance were compared using Welch’s *t* test. The means of nonnormally distributed data were compared using a nonparametric Wilcoxon text with post hoc pairwise Wilcoxon tests. All statistical analyses were performed using JMP Pro 17.0 software (SAS Institute Inc., Cary, NC, USA). *P* values of less than 0.05 were considered statistically significant.

## Acknowledgements

*Gardnerella* strains 3336 and 2492 were provided by Dr. Marilyn Roberts. Strains ATCC 14018, AMD, 101, JCP7275, and UM035 were provided by Dr. Kim Jefferson. Strains 1400E, 1500E, and 061/19V5 were provided by Dr. Sharon Hillier. Strain Gv1, the parent strain of AKK101 was provided by Dr. Caitlin Pepperell. *L. gasseri* 33323 was provided by Dr. Cindy Arvidson. Proteomics (enzymatic digestion through analysis) was performed by the UW-Madison Biotechnology Center Mass Spectrometry Core. Whole genome sequencing was performed by the Virginia Commonwealth University Genomics Core.

---

**Figure S1.**
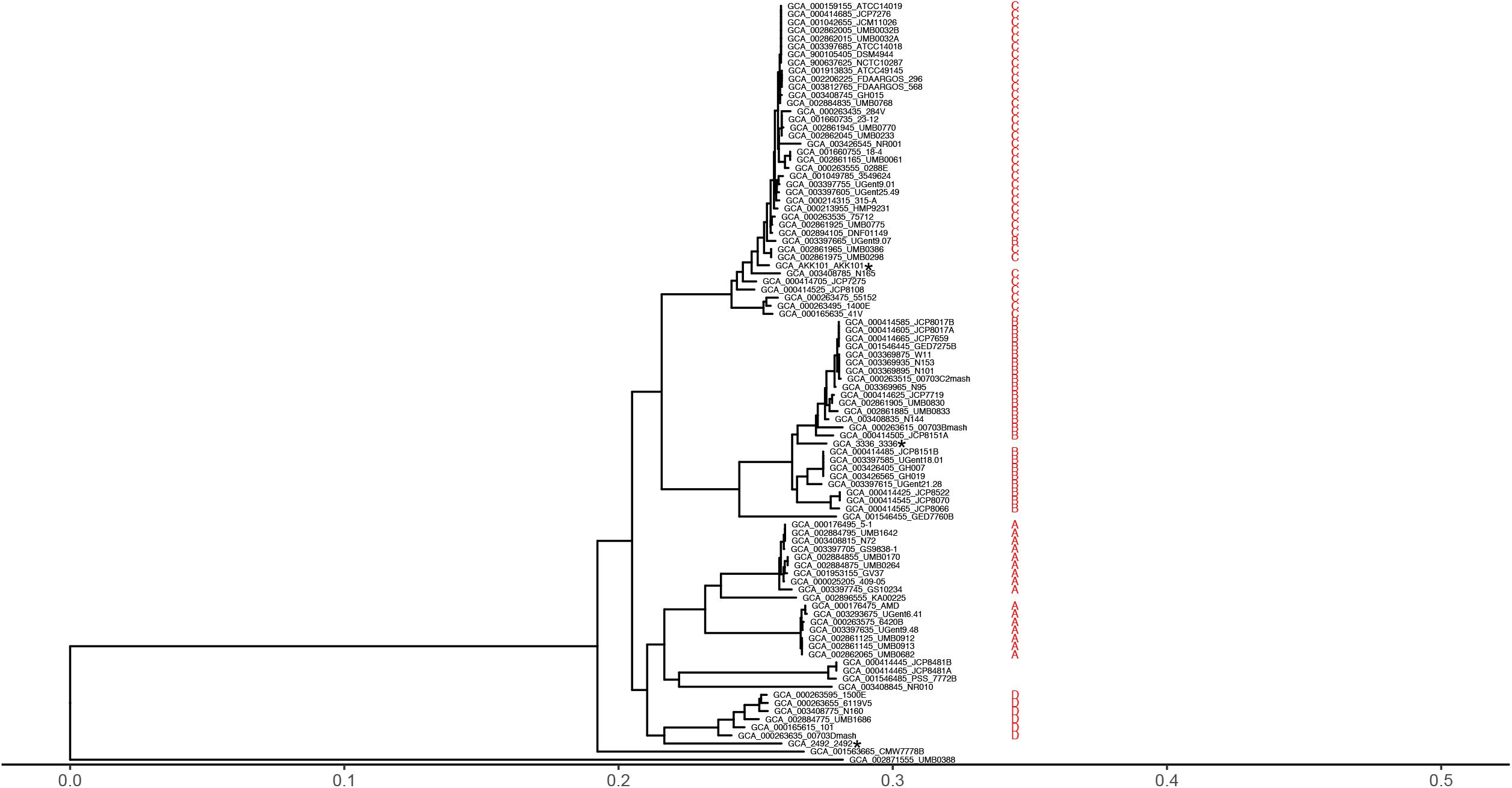
*cpn60* alignment to determine species designation of *Gardnerella* strains of interest. Neighbor-joining phylogenetic tree of full length *cpn60* nucleotide sequences (1629 bp) of 91 *Gardnerella* spp. available genomes from GenBank. Clade distribution as described by Jayaprakash et al. 2012. AKK101 most closely groups with clade C and strain 3336 with clade B. Strain 2492 is closest to clade D, but may be outside of the original 4 clade designation. Regardless, each strain of interest is a distinct *Gardnerella* species. * Marks strains characterized in this study.

**Table S1.**
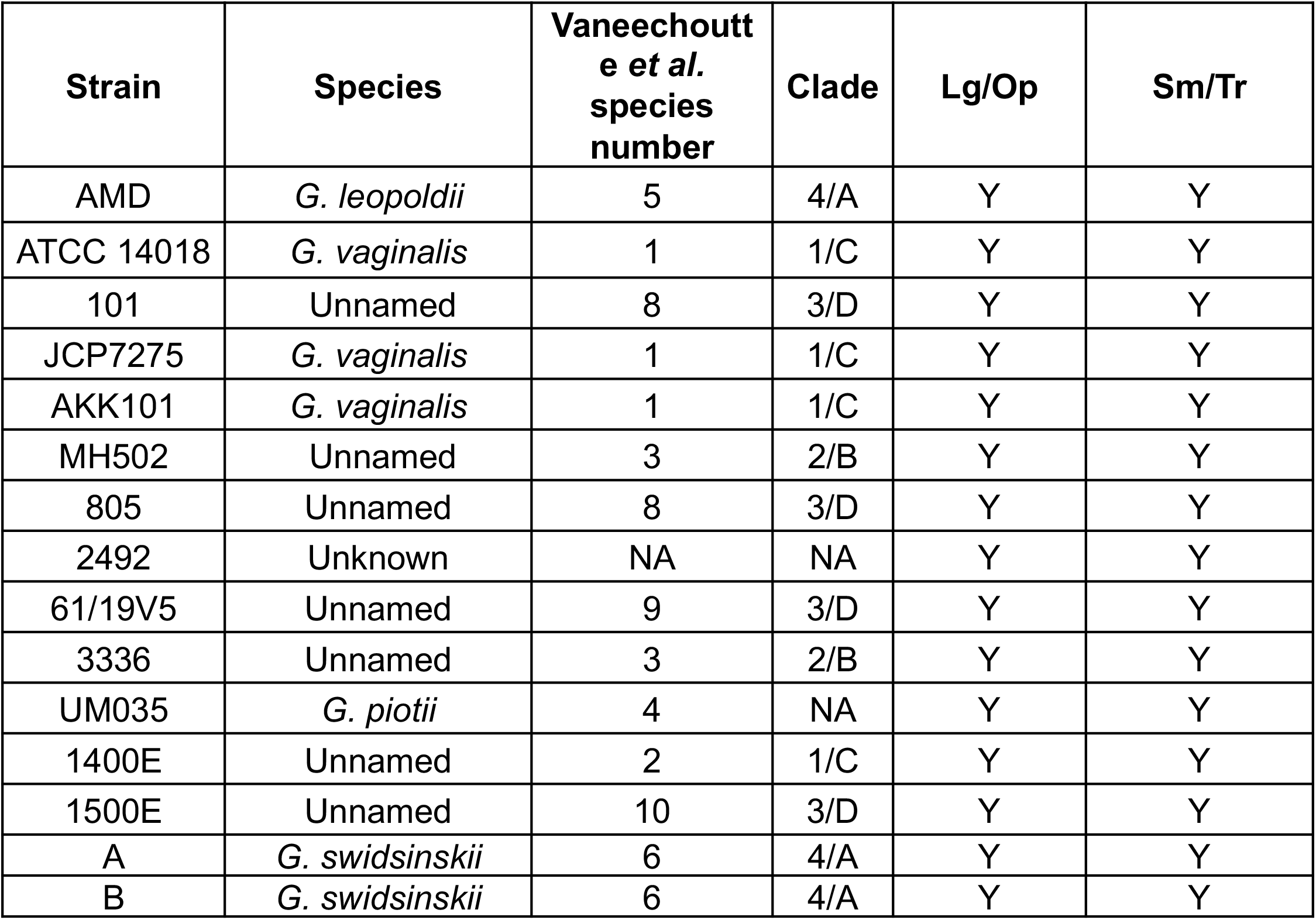
Various *Gardnerella* species exhibit both colony phenotypes. The presence of both phenotypes was confirmed by streaking strains onto BHIFF (colony size) or sBHI agar plates supplemented with 10% FBS (opacity). Only four of the proposed thirteen Vaneechoutte *Gardnerella* species have been given a species name designation. The remaining nine are only recognized by their number. “NA” means that a strain cannot be classified into one of the 13 Vaneechoutte species groups. These species designations were determined by *cpn60* sequence alignment (Fig. S1). “Y” = yes, both phenotypes observed.

**Figure S2.**
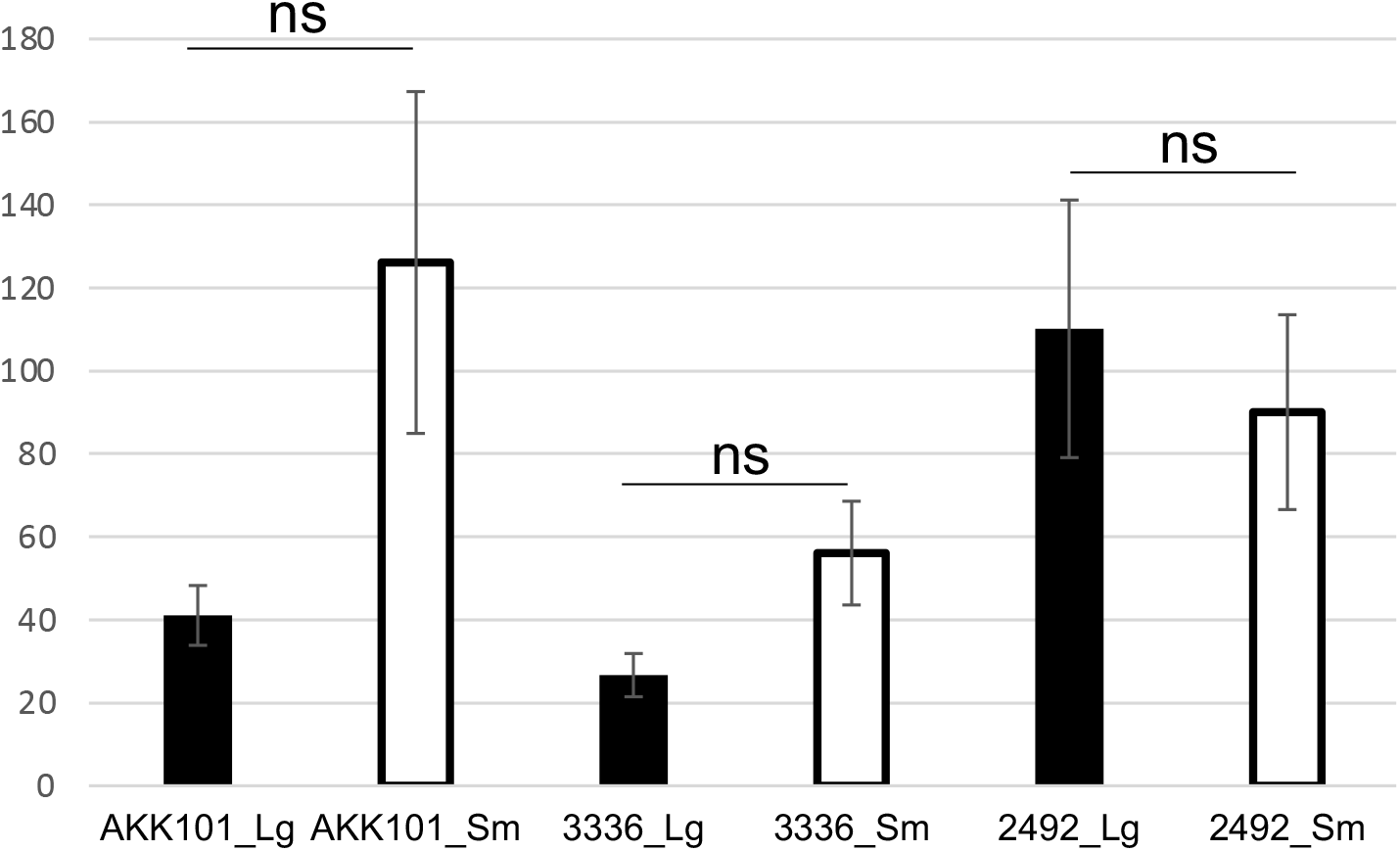
*Gardnerella* variant doubling times in biofilm medium. Doubling time for each strain and variant in biofilm medium. Error bars represent the standard error of the mean between at least three independent experiments. “ns” = not significant.

**Figure S3.**
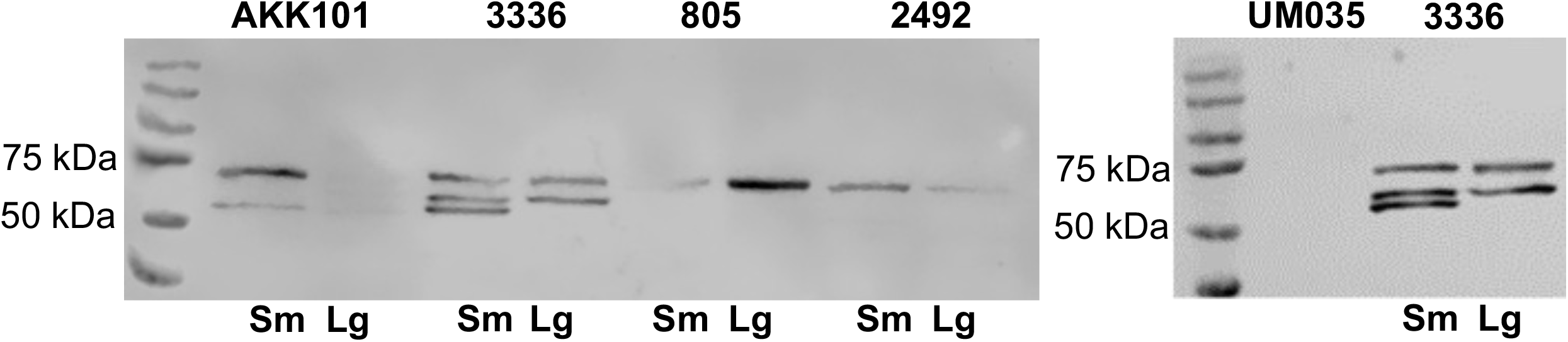
VLY Western blots of *Gardnerella* spp. supernatants. Strain 805 was not characterized in this study but was included for continuity of the blot. Strain UM035 was previously reported as *vly* deficient (Castro et al. 2015). It is unclear why multiple bands are present for some strains.

**Figure S4.**
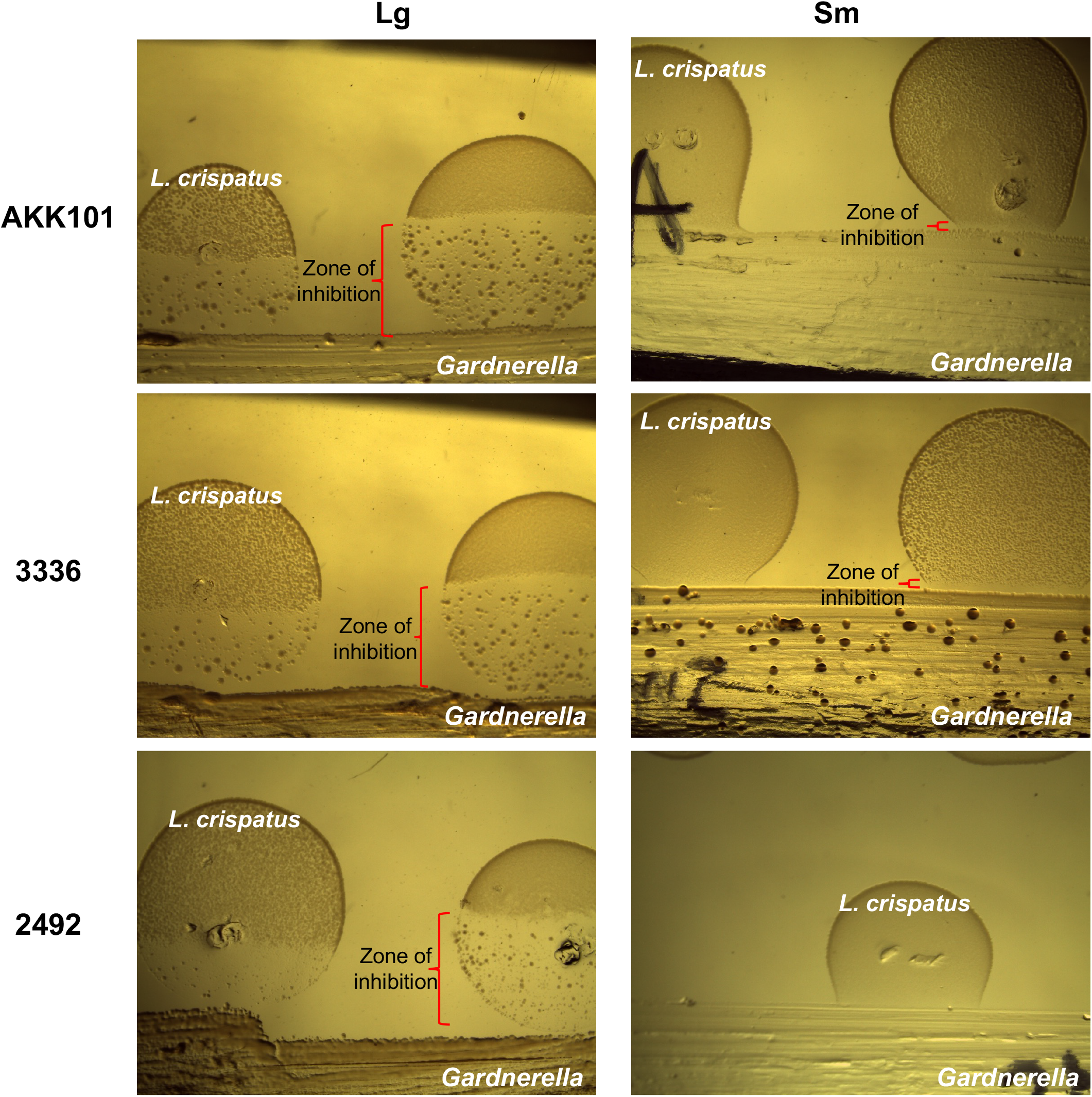
Colony variants differ in their antagonism of *L. crispatus*. *Gardnerella* colony variants were streaked onto biofilm medium + 10% FBS and grown for 24 hours before traversing the streak with 5µl of diluted *L. crispatus* cultures. Results from a representative experiment are shown but three independent experiments were performed. Images were taken on a Leica S8 APO stereo microscope.

**Table S2.** Proteomic differences between opaque and translucent colony variants. The top 15 proteins with highest Lg/Sm ratio (fold difference) in strain 2492 are shown. Data are presented as an average of three independent experiments.

| Protein annotation | Gene name | Fold difference |
| --- | --- | --- |
| Putative calcium-translocating P-type ATPase, PMCA-type | HMPREF0421_20193 | 18.5 |
| DNA topoisomerase (ATP-hydrolyzing) | HMPREF0421_20607 | 15.5 |
| Aminotransferase, class I/II | HMPREF0421_20226 | 4.9 |
| Ribonuclease J | *rnj* | 4.8 |
| 10 kDa chaperonin | *groS* | 4.4 |
| Peptidase C1-like family | HMPREF0421_20285 | 3.4 |
| Pyridine nucleotide-disulfide oxidoreductase | HMPREF0421_20163 | 3.4 |
| Glutamine synthetase | *glnA* | 3.1 |
| ABC transporter, substrate-binding protein | HMPREF0421_20686 | 2.8 |
| Tyrosine--tRNA ligase | *tyrS* | 2.7 |
| GTP-binding protein TypA | *typA* | 2.5 |
| 60 kDa chaperonin | *groL* | 2.45 |
| Ribonucleoside-diphosphate reductase | HMPREF0421_21332 | 2.45 |
| Ribose-phosphate pyrophosphokinase | *prs* | 2.5 |
| DNA-directed RNA polymerase subunit alpha | *rpoA* | 2.4 |

**Table S3.** Proteomic differences between opaque and translucent colony variants. The top 15 proteins with highest Lg/Sm ratio (fold difference) in strain ATCC 14018 are shown. Data are presented as an average of three independent experiments.

| Protein annotation | Gene name | Fold difference |
| --- | --- | --- |
| Repeat protein | HMPREF0421_21350 | 43 |
| Cysteine synthase | *cysK* | 12 |
| Alpha-L-fucosidase | HMPREF0421_20101 | 9.8 |
| ATP synthase, delta/epsilon subunit, beta-sandwich domain protein | HMPREF0421_20002 | 9.2 |
| ATP synthase subunit b | *atpF* | 8.4 |
| ABC transporter, solute-binding protein | HMPREF0421_20097 | 8.3 |
| DnaJ domain protein | HMPREF0421_20259 | 6.8 |
| Serine acetyltransferase | HMPREF0421_21032 | 5.6 |
| Ribokinase | *rbsK* | 5.4 |
| Argininosuccinate synthase | *argG* | 5.3 |
| ABC transporter, solute-binding protein | HMPREF0421_20451 | 5.2 |
| Aminotransferase, class I/II | HMPREF0421_20226 | 5 |
| ABC transporter, ATP-binding protein | HMPREF0421_21100 | 4.9 |
| Beta-galactosidase | HMPREF0421_20100 | 4.7 |
| SUF system FeS assembly protein, NifU family | HMPREF0421_20529 | 3.6 |

**Table S4.** Proteomic differences between translucent and opaque colony variants. The top 15 proteins with highest Sm/Lg ratio (fold difference) in strain 2492 are shown. Data are presented as an average of three independent experiments.

| Protein annotation | Gene name | Fold difference |
| --- | --- | --- |
| Thiol-activated cytolysin | HMPREF0421_20066 | 95 |
| 4-alpha-glucanotransferase | *malQ* | 23.7 |
| Phosphomethylpyrimidine synthase | *thiC* | 15 |
| ABC transporter, ATP-binding protein | HMPREF0421_20038 | 11 |
| Ribosome hibernation promoting factor | *raiA* | 9.9 |
| NADH oxidase | *noxE* | 8 |
| 3-deoxy-7-phosphoheptulonate synthase | HMPREF0421_20347 | 8 |
| Hydrolase, alpha/beta domain protein | HMPREF0421_20703 | 7.3 |
| GA module protein | HMPREF0421_21196 | 7 |
| Oxidoreductase, aldo/keto reductase family protein | HMPREF0421_20336 | 5.3 |
| Formate C-acetyltransferase | *pflB* | 5.1 |
| ATPase family associated with various cellular activities (AAA) | HMPREF0421_20653 | 5 |
| Universal stress family protein | HMPREF0421_20544 | 4.7 |
| 2-dehydropantoate 2-reductase | HMPREF0421_20203 | 4.6 |
| ROK family protein | HMPREF0421_20797 | 4.5 |

**Table S5.**
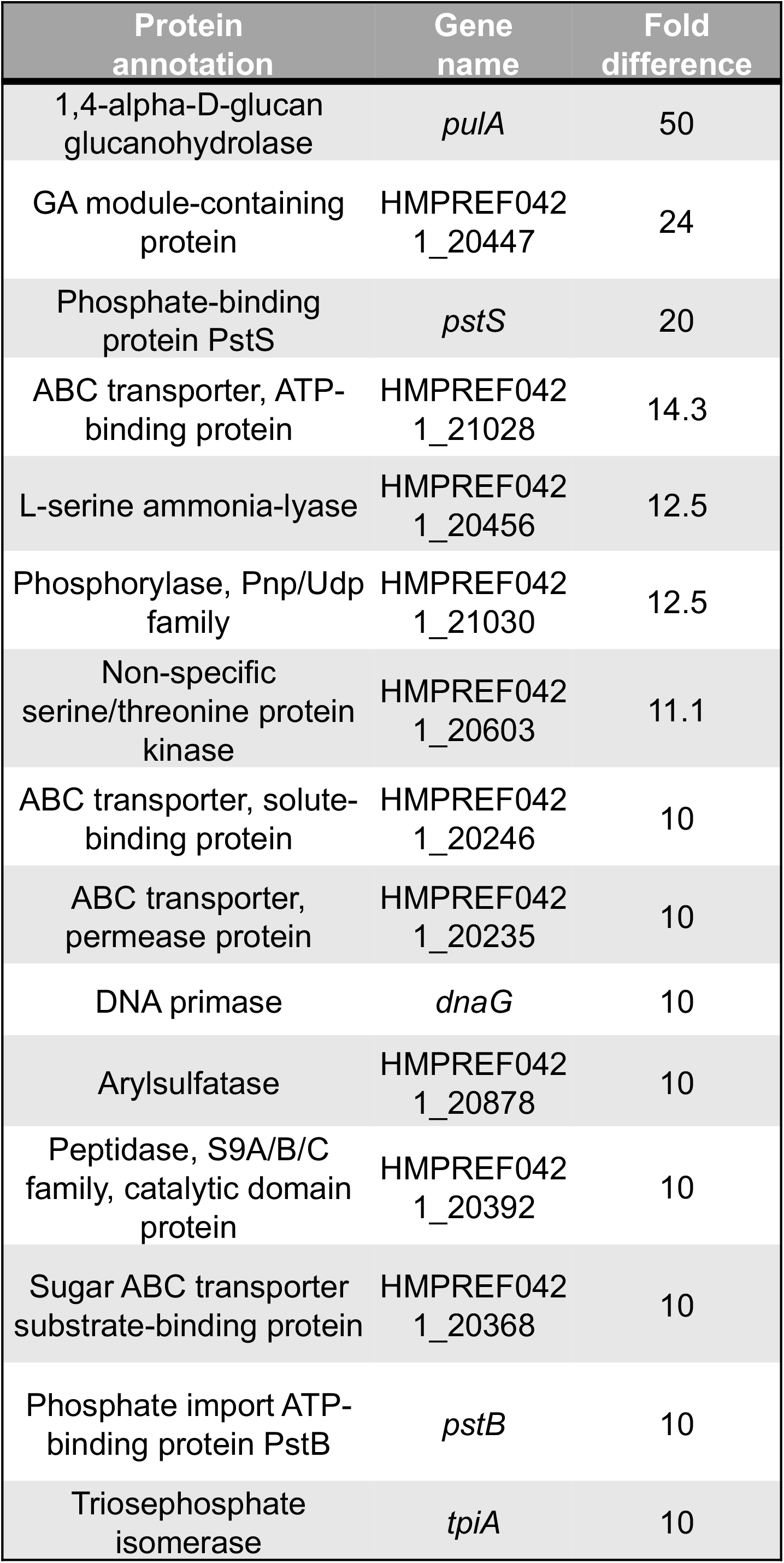
Proteomic differences between translucent and opaque colony variants. The top 15 proteins with highest Sm/Lg ratio (fold difference) in strain ATCC 14018 are shown. Data are presented as an average of three independent experiments.

**Figure S3.**
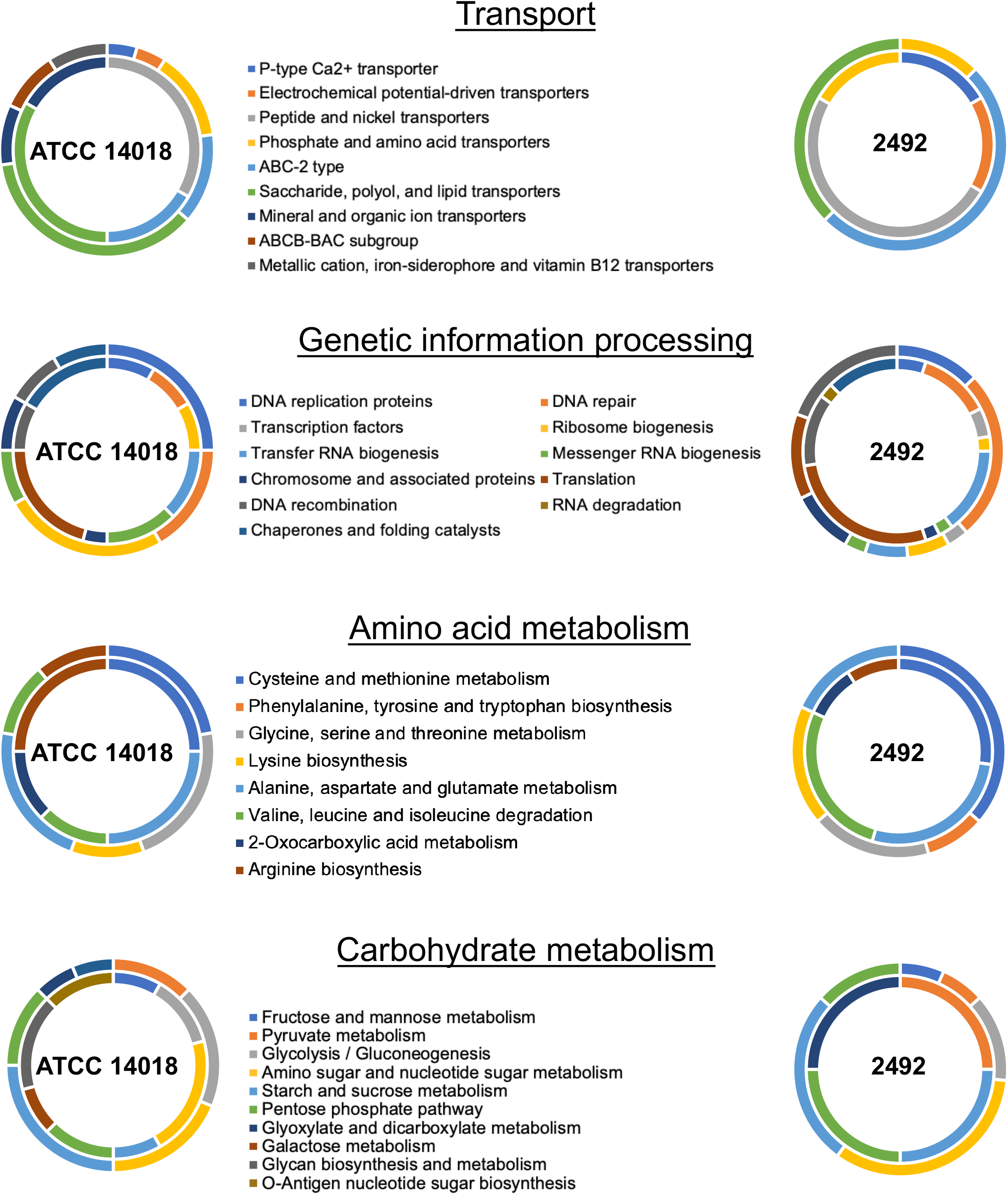
KEGG protein subclasses. Inner ring is the Lg variant, outer ring is the Sm variant.

## Notes

### Competing Interest Statement

The authors have declared no competing interest.

## References

1. Catlin, B. W. Gardnerella vaginalis: characteristics, clinical considerations, and controversies. Clin Microbiol Rev 5, 213–37 (1992).

2. Muzny, C. A. & Schwebke, J. R. Gardnerella vaginalis: Still a prime suspect in the pathogenesis of bacterial vaginosis. Curr Infect Dis Rep 15, 130–135 (2013).

3. Muzny, C. A., Łaniewski, P., Schwebke, J. R. & Herbst-Kralovetz, M. M. Host–vaginal microbiota interactions in the pathogenesis of bacterial vaginosis. Curr Opin Infect Dis 33, 59–65 (2020).

4. Morrill, S. R., Gilbert, N. M. & Lewis, A. L. Gardnerella vaginalis as a direct cause or an indirect mediator of polymicrobial pathogenesis in bacterial vaginosis: Appraisal of the in vivo evidence from experimental models. Front Cell Infect Microbiol 10, 168 (2020).

5. Zevin, A. S. et al. Microbiome Composition and Function Drives Wound-Healing Impairment in the Female Genital Tract. PLoS Pathog 12, e1005889 (2016).

6. Hoang, T. et al. The cervicovaginal mucus barrier to HIV-1 is diminished in bacterial vaginosis. PLoS Pathog 16, e1008236 (2020).

7. Garcia, E. M., Kraskauskiene, V., Koblinski, J. E. & Jefferson, K. K. Interaction of gardnerella vaginalis and vaginolysin with the apical versus basolateral face of a three-dimensional model of vaginal epithelium. Infect Immun 87, (2019).

8. Muzny, C. A., Łaniewski, P., Schwebke, J. R. & Herbst-Kralovetz, M. M. Host–vaginal microbiota interactions in the pathogenesis of bacterial vaginosis. Curr Opin Infect Dis 33, 59–65 (2020).

9. Pleckaityte, M., Janulaitiene, M., Lasickiene, R. & Zvirbliene, A. Genetic and biochemical diversity of Gardnerella vaginalis strains isolated from women with bacterial vaginosis. FEMS Immunol Med Microbiol 65, 69–77 (2012).

10. Ahmed, A. et al. Comparative genomic analyses of 17 clinical isolates of Gardnerella vaginalis provide evidence of multiple genetically isolated clades consistent with subspeciation into genovars. J Bacteriol 194, 3922–3937 (2012).

11. Cornejo, O. E., Hickey, R. J., Suzuki, H. & Forney, L. J. Focusing the diversity of Gardnerella vaginalis through the lens of ecotypes. Evol Appl 11, 312–324 (2018).

12. Yeoman, C. J. et al. Comparative genomics of Gardnerella vaginalis strains reveals substantial differences in metabolic and virulence potential. PLoS One 5, e12411 (2010).

13. Hummelen, R. et al. Deep sequencing of the vaginal microbiota of women with HIV. PLoS One 5, e12078 (2010).

14. Hill, J. E. et al. Characterization of vaginal microflora of healthy, nonpregnant women by chaperonin-60 sequence-based methods. Am J Obstet Gynecol 193, 682–692 (2005).

15. Janulaitiene, M. et al. Phenotypic characterization of Gardnerella vaginalis subgroups suggests differences in their virulence potential. PLoS One 13, (2018).

16. Vaneechoutte, M. et al. Emended description of Gardnerella vaginalis and description of gardnerella leopoldii sp. Nov., gardnerella piotii sp. nov. and Gardnerella swidsinskii sp. nov., with delineation of 13 genomic species within the genus Gardnerella. Int J Syst Evol Microbiol 69, 679–687 (2019).

17. Balashov, S. V., Mordechai, E., Adelson, M. E. & Gygax, S. E. Identification, quantification and subtyping of Gardnerella vaginalis in noncultured clinical vaginal samples by quantitative PCR. J Med Microbiol 63, 162–175 (2014).

18. Garcia, E. M. et al. Sequence comparison of vaginolysin from different gardnerella species. Pathogens 10, 1–15 (2021).

19. Hill, J. E. & Albert, A. Y. K. Resolution and co-occurrence patterns of Gardnerella leopoldii, Gardnerella swidsinskii, Gardnerella piotii and Gardnerella vaginalis within the vaginal microbiome. Infect Immun (2019) doi:10.1128/IAI.00532-19.

20. Paramel Jayaprakash, T., Schellenberg, J. J. & Hill, J. E. Resolution and characterization of distinct cpn60-based subgroups of gardnerella vaginalis in the vaginal microbiota. PLoS One (2012) doi:10.1371/journal.pone.0043009.

21. Janulaitiene, M. et al. Phenotypic characterization of Gardnerella vaginalis subgroups suggests differences in their virulence potential. PLoS One 13, e0200625 (2018).

22. Shipitsyna, E. et al. Quantitation of all Four Gardnerella vaginalis Clades Detects Abnormal Vaginal Microbiota Characteristic of Bacterial Vaginosis More Accurately than Putative G. vaginalis Sialidase A Gene Count. Mol Diagn Ther 23, 139–147 (2019).

23. Abraham, J. M., Freitag, C. S., Clements, J. R. & Eisenstein, B. I. An invertible element of DNA controls phase variation of type 1 fimbriae of Escherichia coli. Proc Natl Acad Sci U S A 82, 5724–5727 (1985).

24. Zhao, H., Li, X., Johnson, D. E., Blomfield, I. & Mobley, H. L. T. In vivo phase variation of MR/P fimbrial gene expression in Proteus mirabilis infecting the urinary tract. Mol Microbiol 23, 1009–1019 (1997).

25. Willems, R., Paul, A., van der Heide, H. G., ter Avest, A. R. & Mooi, F. R. Fimbrial phase variation in Bordetella pertussis: a novel mechanism for transcriptional regulation. EMBO J 9, 2803–9 (1990).

26. Nicholson, B. & Low, D. DNA methylation-dependent regulation of Pef expression in Salmonella typhimurium. Mol Microbiol 35, 728–742 (2000).

27. Ham, S. M., Alphen, L., Mooi, F. R. & Van Putten”, J. P. M. Phase Variation of H. influenzae Fimbriae: Transcriptional Control of Two Divergent Genes through a Variable Combined Promoter Region. Cell vol. 73 (1993).

28. Henderson, I. R., Owen, P. & Nataro, J. P. Molecular switches - the ON and OFF of bacterial phase variation. Mol Microbiol 33, 919–932 (1999).

29. Moxon, R., Bayliss, C. & Hood, D. Bacterial contingency loci: the role of simple sequence DNA repeats in bacterial adaptation. Annu Rev Genet 40, 307–333 (2006).

30. Meyer, T. F., Gibbs, C. P. & Haas, R. VARIATION AND CONTROL OF PROTEIN EXPRESSION IN NEISSERIA. https://doi.org/10.1146/annurev.mi.44.100190.002315 44, p451–477 (2003).

31. Alcott, A. M. et al. Variable Expression of Opa Proteins by Neisseria gonorrhoeae Influences Bacterial Association and Phagocytic Killing by Human Neutrophils. J Bacteriol 204, (2022).

32. Sintsova, A. et al. Selection for a CEACAM receptor-specific binding phenotype during Neisseria gonorrhoeae infection of the human genital tract. Infect Immun 83, 1372–1383 (2015).

33. Grant, C. C. R., Bos, M. P. & Belland, R. J. Proteoglycan receptor binding by Neisseria gonorrhoeae MS11 is determined by the HV-1 region of OpaA. Mol Microbiol 32, 233–242 (1999).

34. Swanson, J. Colony opacity and protein II compositions of gonococci. Infect Immun 37, 359–68 (1982).

35. Swanson, J. & Barrera, O. Gonococcal pilus subunit size heterogeneity correlates with transitions in colony piliation phenotype, not with changes in colony opacity. Journal of Experimental Medicine 158, 1459–1472 (1983).

36. Inouye, T., Ohta, H., Kokeguchi, S., Fukui, K. & Kato, K. Colonial variation and fimbriation of Actinobacillus actinomycetemcomitans. FEMS Microbiol Lett 69, 13–17 (1990).

37. Dean, M. A., Olsen, R. J., Wesley Long, S., Rosato, A. E. & Musser, J. M. Identification of point mutations in clinical staphylococcus aureus strains that produce small-colony variants auxotrophic for menadione. Infect Immun 82, 1600–1605 (2014).

38. Chee, W. J. Y., Chew, S. Y. & Than, L. T. L. Vaginal microbiota and the potential of Lactobacillus derivatives in maintaining vaginal health. Microbial Cell Factories 2020 19:1 19, 1–24 (2020).

39. Muzny, C. A. et al. An Updated Conceptual Model on the Pathogenesis of Bacterial Vaginosis. J Infect Dis 220, 1399 (2019).

40. Teixeira, G. S. et al. Characteristics of Lactobacillus and Gardnerella vaginalis from women with or without bacterial vaginosis and their relationships in gnotobiotic mice. J Med Microbiol 61, 1074–1081 (2012).

41. van der Woude, M. W. & Bäumler, A. J. Phase and Antigenic Variation in Bacteria. Clin Microbiol Rev 17, 581 (2004).

42. Kurukulasuriya, S. P., Patterson, M. H. & Hill, J. E. Slipped-strand mispairing in the gene encoding sialidase NanH3 in gardnerella spp. Infect Immun 89, (2021).

43. Robinson, L. S., Schwebke, J., Lewis, W. G. & Lewis, A. L. Identification and characterization of NanH2 and NanH3, enzymes responsible for sialidase activity in the vaginal bacterium Gardnerella vaginalis. Journal of Biological Chemistry 294, 5230–5245 (2019).

44. Dautry-Varsat, A., Subtil, A. & Hackstadt, T. Recent insights into the mechanisms of Chlamydia entry. Cell Microbiol 7, 1714–1722 (2005).

45. Kvint, K., Nachin, L., Diez, A. & Nyström, T. The bacterial universal stress protein: function and regulation. Curr Opin Microbiol 6, 140–145 (2003).

46. Ueta, M. et al. Role of HPF (hibernation promoting factor) in translational activity in Escherichia coli. J Biochem 143, 425–433 (2008).

47. Sadarangani, M., Pollard, A. J. & Gray-Owen, S. D. Opa proteins and CEACAMs: pathways of immune engagement for pathogenic Neisseria. FEMS Microbiol Rev 35, 498–514 (2011).

48. Russell, M. W., Jerse, A. E. & Gray-Owen, S. D. Progress Toward a Gonococcal Vaccine: The Way Forward. Front Immunol 10, 2417 (2019).

49. Proctor, R. A. et al. Small colony variants: a pathogenic form of bacteria that facilitates persistent and recurrent infections. Nature Reviews Microbiology 2006 4:4 4, 295–305 (2006).

50. Kellogg, D. S., Peacock, W. L., Deacon, W. E., Brown, L. & Pirkle, C. I. Neisseria gonorrhoeae Virulence genetically linked to clonal variation. Infectious Diseases in Obstetrics and Gynecology, Sixth Edition 191–201 (2008) doi:10.1128/JB.85.6.1274-1279.1963.

51. Seemann, T. Prokka: rapid prokaryotic genome annotation. Bioinformatics 30, 2068–2069 (2014).

52. Yu, G., Smith, D. K., Zhu, H., Guan, Y. & Lam, T. T. Y. ggtree: an r package for visualization and annotation of phylogenetic trees with their covariates and other associated data. Methods Ecol Evol 8, 28–36 (2017).

53. Dereeper, A. et al. Phylogeny.fr: robust phylogenetic analysis for the non-specialist. Nucleic Acids Res 36, (2008).

54. Paramel Jayaprakash, T., Schellenberg, J. J. & Hill, J. E. Resolution and Characterization of Distinct cpn60-Based Subgroups of Gardnerella vaginalis in the Vaginal Microbiota. PLoS One 7, e43009 (2012).

55. Vaneechoutte, M. et al. Emended description of Gardnerella vaginalis and description of gardnerella leopoldii sp. Nov., gardnerella piotii sp. nov. and Gardnerella swidsinskii sp. nov., with delineation of 13 genomic species within the genus Gardnerella. Int J Syst Evol Microbiol 69, 679–687 (2019).

56. Ahmed, A. et al. Comparative genomic analyses of 17 clinical isolates of Gardnerella vaginalis provide evidence of multiple genetically isolated clades consistent with subspeciation into genovars. J Bacteriol 194, 3922–3937 (2012).

57. Ramsey, M. E. et al. TraK and traB are conserved outer membrane proteins of the Neisseria gonorrhoeae type IV secretion system and are expressed at low levels in wild-type cells. J Bacteriol 196, 2954–2968 (2014).

58. Pleckaityte, M., Mistiniene, E., Lasickiene, R., Zvirblis, G. & Zvirbliene, A. Generation of recombinant single-chain antibodies neutralizing the cytolytic activity of vaginolysin, the main virulence factor of Gardnerella vaginalis. BMC Biotechnol 11, 100 (2011).

59. Castro, J. et al. Using an in-vitro biofilm model to assess the virulence potential of Bacterial Vaginosis or non-Bacterial Vaginosis Gardnerella vaginalis isolates. Scientific Reports 2015 5:1 5, 1–10 (2015).

60. Chambers, M. C. et al. A cross-platform toolkit for mass spectrometry and proteomics. Nature Biotechnology 2012 30:10 30, 918–920 (2012).

61. Nesvizhskii, A. I., Keller, A., Kolker, E. & Aebersold, R. A statistical model for identifying proteins by tandem mass spectrometry. Anal Chem 75, 4646–4658 (2003).

62. Kanehisa, M., Sato, Y., Kawashima, M., Furumichi, M. & Tanabe, M. KEGG as a reference resource for gene and protein annotation. Nucleic Acids Res 44, D457–D462 (2016).

63. Kolmogorov, M., Yuan, J., Lin, Y. & Pevzner, P. A. Assembly of long, error-prone reads using repeat graphs. Nature Biotechnology 2019 37:5 37, 540–546 (2019).

64. Garrison, E. & Marth, G. Haplotype-based variant detection from short-read sequencing. (2012) doi:10.48550/arxiv.1207.3907.

65. Cingolani, P. et al. A program for annotating and predicting the effects of single nucleotide polymorphisms, SnpEff: SNPs in the genome of Drosophila melanogaster strain w1118; iso-2; iso-3. Fly (Austin) 6, 80 (2012).

